# Pan-genome analysis of different morphotypes reveals genomic basis of *Brassica oleracea* domestication and differential organogenesis

**DOI:** 10.1101/2023.10.24.563711

**Authors:** Ning Guo, Shenyun Wang, Tianyi Wang, Mengmeng Duan, Mei Zong, Liming Miao, Shuo Han, Guixiang Wang, Xin Liu, Deshuang Zhang, Chengzhi Jiao, Hongwei Xu, Liyang Chen, Zhangjun Fei, Jianbin Li, Fan Liu

**Affiliations:** State Key Laboratory of Vegetable Biobreeding, National Engineering Research Center for Vegetables, Beijing Key Laboratory of Vegetable Germplasms Improvement, Key Laboratory of Biology and Genetics Improvement of Horticultural Crops (North China), Beijing Vegetable Research Center, Beijing Academy of Agriculture and Forestry Science, Beijing 100097, China; Jiangsu Key Laboratory for Horticultural Crop Genetic Improvement, Vegetable Research Institute, Jiangsu Academy of Agricultural Science, Nanjing, Jiangsu, China; Smartgenomics Technology Institute, Tianjin 301700, China; Boyce Thompson Institute, Ithaca, NY, USA

## Abstract

The domestication of *Brassica oleracea* has resulted in diverse morphological types with distinct patterns of organ development. Here we report a graph-based pan-genome of *B. oleracea* constructed with high-quality genome assemblies of different morphotypes. The pan-genome harbors over 200 structural variant (SV) hotspot regions enriched with auxin and flowering-related genes. Population genomic analyses reveal that early domestication of *B. oleracea* focused on leaf or stem selection. Gene flows resulting from agricultural practices and variety improvement are detected among different morphotypes. Selective sweep analysis identifies an auxin-responsive SAUR gene and a CLE family gene as the crucial players in the leaf-stem differentiation during the early stage of *B. oleracea* domestication, and the *BoKAN1* gene as instrumental in shaping the leafy heads of cabbage and Brussels sprouts. Our pan-genome and functional analyses further discover that variations in the *BoFLC2* gene play key roles in the divergence of vernalization and flowering characteristics among different morphotypes, and variations in the first intron of *BoFLC3* are involved in fine-tuning the flowering process in cauliflower. This study provides a comprehensive understanding of the pan-genome of *B. oleracea* and sheds light on the domestication and differential organ development of this globally important crop species.

## Introduction

*Brassica oleracea*, a member of the Brassicaceae family, is an economically important vegetable crop cultivated worldwide and encompasses a remarkable diversity of morphotypes. Domestication and subsequent breeding of *B. oleracea* have played a crucial role in shaping its morphological diversity, resulting in the emergence of numerous cultivars with varying appearances and agronomic traits (Cheng et al., 2016), such as leafy heads, curds, tuberous stems, and flower stalks. Additionally, selections aimed to improve plant adaptation to diverse geographic regions and seasons have enriched the diversity of flowering characteristics in *B. oleracea* (Maggioni, 2015). Gaining a comprehensive understanding of the genetic mechanisms underlying these phenotypic variations holds significant importance for crop enhancement and the long-term sustainability of agriculture.

Traditional genetic and recent genomic studies have shed light on the genetic control of specific traits in *B. oleracea* (Arias et al., 2021; Cheng et al., 2016; Guo et al., 2021; Ji et al., 2023). However, to fully understand the genomic variations responsible for the morphological diversity in *B. oleracea*, it is crucial to conduct comprehensive investigations into the pan-genome as a single reference genome may result in the loss of a significant amount of meaningful genetic information (Bayer et al., 2020; Danilevicz et al., 2020; Lei et al., 2021; Rehman et al., 2022; Wang et al., 2023b). In recent years, advances in sequencing technologies and bioinformatics have made pan-genome analysis feasible and affordable. The availability of multiple high-quality reference genomes and the emergence of powerful computational tools have enabled the construction and comparison of pan-genomes for various plant species (Cai et al., 2021; Li et al., 2023; Liu et al., 2020; Qin et al., 2021; Wang et al., 2023a; Wang et al., 2022b; Zhang et al., 2021). These pan-genomic investigations have provided valuable insights into the evolutionary dynamics, genetic diversity, and functional variation within/among species, highlighting the importance of studying the pan-genome for a comprehensive understanding of plant biology. Previous pan-genome investigation of *B. oleracea*, which employed Illumina short-read resequencing and a reference-guided approach, did not utilize robust *de novo* assemblies across diverse morphotypes to create a graph-based pan-genome integrating whole-genome structural variations (Golicz et al., 2016). Consequently, a comprehensive view of species diversification-driving structural variations is lacking. Hence, it is imperative to generate high-quality reference genomes and establish a graph-based pan-genome to fully explore the genetic diversity among different *B. oleracea* morphotypes.

In this study, we first assemble six reference-grade genomes for the wild cabbage and different *B. oleracea* morphotypes, and construct both gene family-based and graph-based pan-genomes from these six and additional five previously published high-quality genome assemblies. Pan-genome analysis identifies structural variant (SV) hotspot regions associated with important agronomical traits. Leveraging the graph-based pan-genome and population analyses, we uncover valuable insights into the evolution and domestication of different *B. oleracea* morphotypes and identify pivotal genes and variations that underlie the differentiation between stem- and leaf-selected morphotypes, the formation of leafy heads, and the divergence of vernalization and flowering traits. The findings from our pan-genome analysis contribute to the understanding of the genomic basis of domestication and organogenesis in *B. oleracea*. Moreover, the pan-genome and genomic variation resources developed in this study provide a foundation for facilitating future biological investigations and the breeding efforts targeting this globally significant crop.

## Results

### High-quality assemblies of six *B. oleracea* morphotypes

We *de novo* assembled genomes of six representative *B. oleracea* accessions from wild cabbage and five different morphotypes, including kale, kohlrabi, broccoli, Brussels sprouts, and Chinese kale (Fig. 1A), using PacBio HiFi long reads. A total of 21.5-30.4 Gb HiFi sequences were generated for each accession, covering about 36.2-49.1× of the six genomes (Supplementary Table 1). The six assemblies had an average length of 608.8 Mb (588.3-634.4 Mb) and an average contig N50 of 13.7 Mb (9.5-17.5 Mb). We then obtained pseudo-chromosomes by anchoring the assembled contigs using a reference-guided approach (Alonge et al., 2019), resulting in an average of 99.5% of contigs being anchored to chromosomes (Supplementary Table 2).

**Fig. 1.**
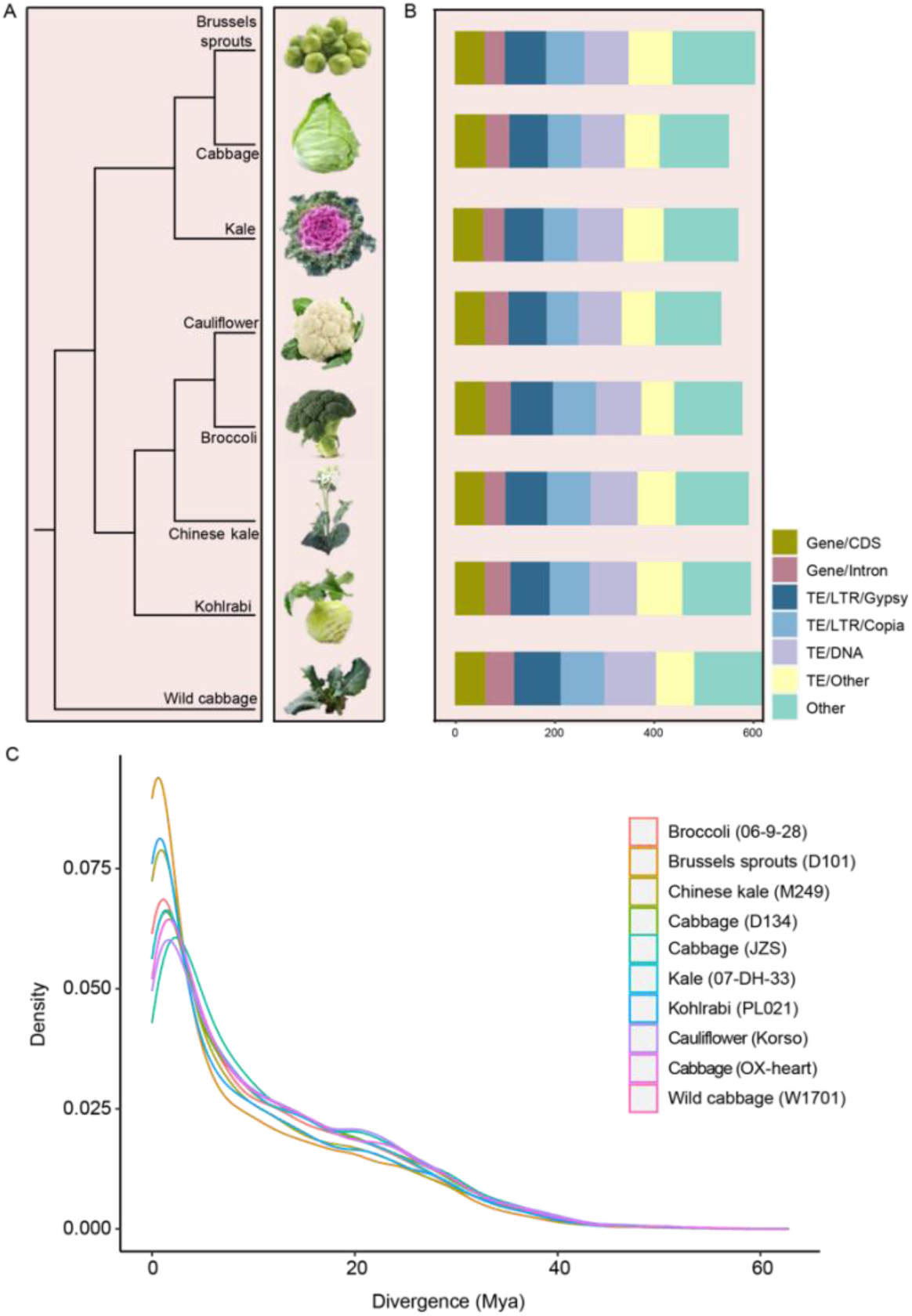
Genome assemblies of different *B. oleracea* morphotypes and the wild cabbage. (A) Phylogenetic relationships of different *B. oleracea* morphotypes and the wild cabbage. The tree was constructed using single-copy genes identified from their genomes. (B) Statistics of *B. oleracea* genome assemblies. Total lengths of each assembly, coding sequences (CDS), introns, Gypsy- and Copia-type long terminal repeats (LTRs), DNA transposable elements (TEs) and other types of TEs are presented. (C) Estimated insertion times of intact LTR retrotransposons in different *B. oleracea* genomes.

BUSCO (Simão et al., 2015) evaluation revealed an average of 99.4% (99.3-99.5%) completeness of the assemblies (Supplementary Table 2). Moreover, the completeness and accuracy of each assembly were further evaluated by mapping ∼50× short reads to each corresponding assembly, revealing a mapping rate of 99.77-99.95% with a very low base error rate (Supplementary Table 3). Furthermore, the LTR Assembly Index (LAI) (Ou et al., 2018) scores for all six assemblies exceeded 10 (Supplementary Table 2). Collectively, the six assemblies were comparable to, or better than, the recently assembled *B. oleracea* genomes with the long reads. Together, these results highlighted the high quality of the six new assemblies of *B. oleracea*.

We predicted an average of 58,750 protein-coding genes for each of the six assemblies. BUSCO evaluation (Simão et al., 2015) of the protein-coding genes showed an average completeness rate of 98.6% (Supplementary Table 2), indicating the high quality of the gene prediction. Repetitive sequences made up ∼57.2% of each genome (55.9-58.6%) (Fig. 1B and Supplementary Table 4). Among the repetitive sequences, long terminal repeat retrotransposons (LTR-RTs) were the most abundant, consistent with previously reported *B. oleracea* genomes (Guo et al., 2021). We further extracted full-length LTR-RTs from the six assemblies, as well as the four previously reported *B. oleracea* genomes (Cai et al., 2020; Guo et al., 2021; Lv et al., 2020). Insertion time estimation of these intact LTR-RTs demonstrated that most of the LTR-RTs were formed recently, less than 2 million years ago (mya) (Fig. 1C and Supplementary Fig. 1), consistent with our previous finding (Guo et al., 2021).

### Gene family-based pan-genome construction

In order to build a gene family-based pan-genome, we reconducted gene predictions of the six new assemblies, and the five recently reported *B. oleracea* genomes [cauliflower (Korso), cabbage (OX-heart, D134, JZS), and broccoli (HDEM)] (Belser et al., 2018; Cai et al., 2020; Guo et al., 2021; Lv et al., 2020), using a unified pipeline to minimize the bias introduced by different methods. This resulted in an average of 61,410 high-confidence protein-coding genes predicted from each genome (Supplementary Table 2). BUSCO evaluation of these protein-coding genes revealed an average completeness rate of 98.4% (Supplementary Table 2). Functional annotations of the predicted protein-coding genes also indicated the high quality of the gene predictions (Supplementary Table 5).

We then constructed the gene family-based pan-genome using the 11 *B. oleracea* genomes with a reported strategy (Hirsch et al., 2014; Hu et al., 2017; Liu et al., 2020; Wang et al., 2018; Zhao et al., 2018). Ortholog clustering grouped all genes from the 11 *B. oleracea* genomes into 62,092 families, of which 27,135 (∼43.70%) were present in all 11 accessions and were defined as core genes, 7,115 (∼11.45%) in 10 accessions (softcore genes), 27,226 (∼43.85%) in 2 to 9 accessions (dispensable genes), and 616 (∼0.99%) in only one accession (specific genes) (Fig. 2A-2C and Supplementary Table 6). These gene families corresponded to 33,821-35,682 core genes, 7,137-8,946 softcore genes, 16,659-20,278 dispensable genes, 39-731 specific genes in each genome (Fig. 2D and Supplementary Table 6).

**Fig. 2.**
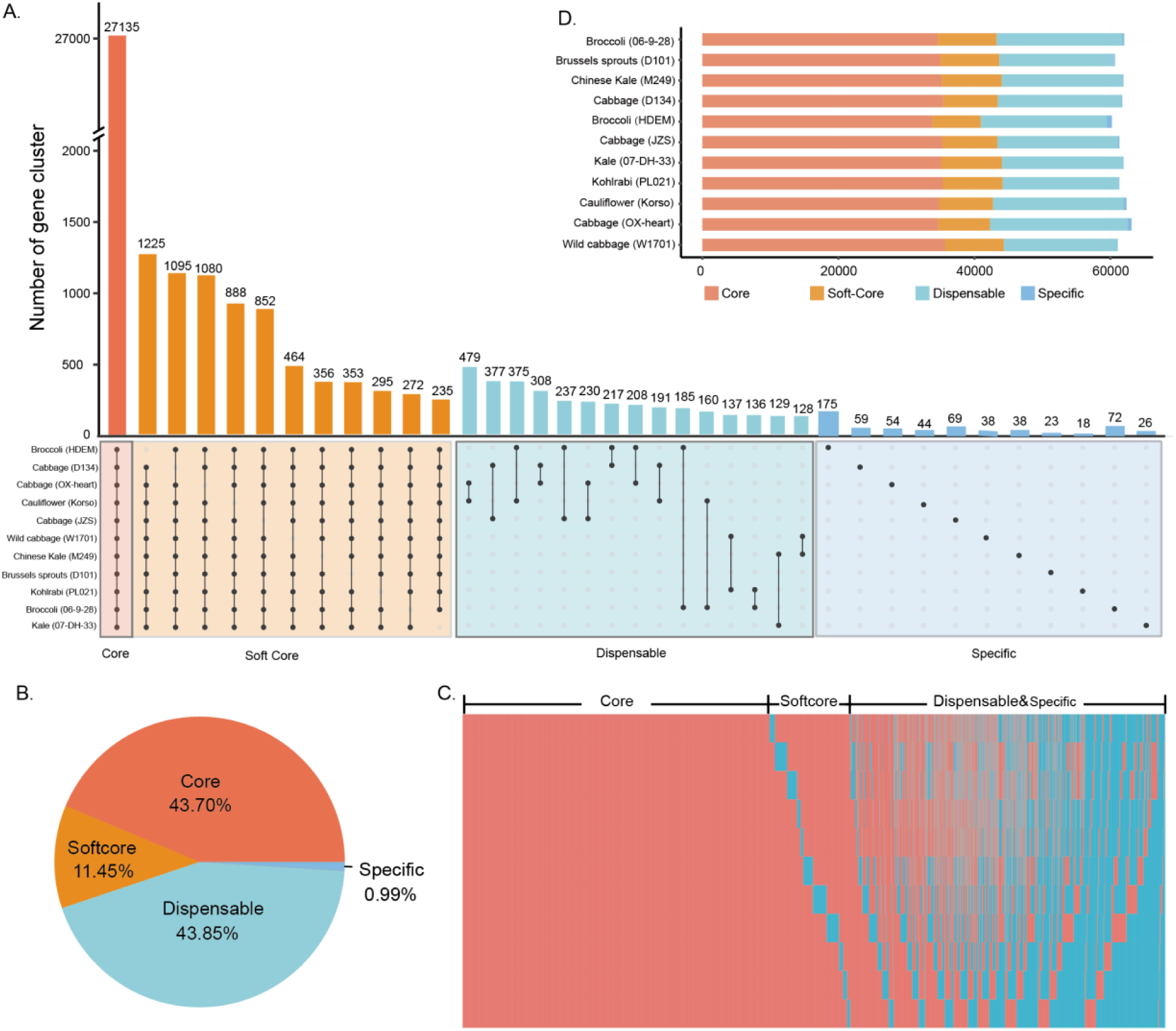
Pan- and core genomes of the 11 *B. oleracea* accessions. (A) UpSet plot of core, softcore, dispensable, and specific gene clusters in the 11 *B. oleracea* genomes. (B) Pie chart showing the proportions of core, softcore, dispensable, and specific gene clusters in the *B. oleracea* pan-genome. (C) Presence (pink) and absence (blue) information of gene families across the 11 *B. oleracea* genomes. (D) Number of core, softcore, dispensable and specific genes in each of the 11 *B. oleracea* genomes.

Gene Ontology (GO) analyses revealed that core genes were predominantly associated with fundamental growth or development processes, and the organization or biogenesis of cellular components, such as protein modification, catabolic processes, chromatin organization, and nitrogen compound transport (Supplementary Fig. 2A). On the other hand, dispensable and private genes were enriched in processes related to environmental response and specific organ genesis, such as response to auxin, cell wall organization, and signal transduction (Supplementary Fig. 2B). This suggests that dispensable and private genes may play significant roles in the adaptation to their specific environments and in their responses to various biotic and abiotic stresses, as well as in their phenotypic diversifications.

### Sequence variation identification and SV hotspots in *B. oleracea* genomes

To discover SVs, we anchored the 10 other genome sequences onto the wild cabbage genome. We identified an average of 159,195 (154,285-170,605) SVs per accession relative to wild cabbage, affecting an average of 170.5Mb (136.2-217.8Mb) of genomic sequences (Fig. 3A-3B and Supplementary Tables 7 and 8). We randomly selected 30 SVs of size ranging from 2 to 8 kb and manually checked them by mapping PacBio long reads to the genome assemblies. We found that all of these SVs were consistent with the breakpoints in the mapping results (Supplementary Fig. 3).

**Fig. 3.**
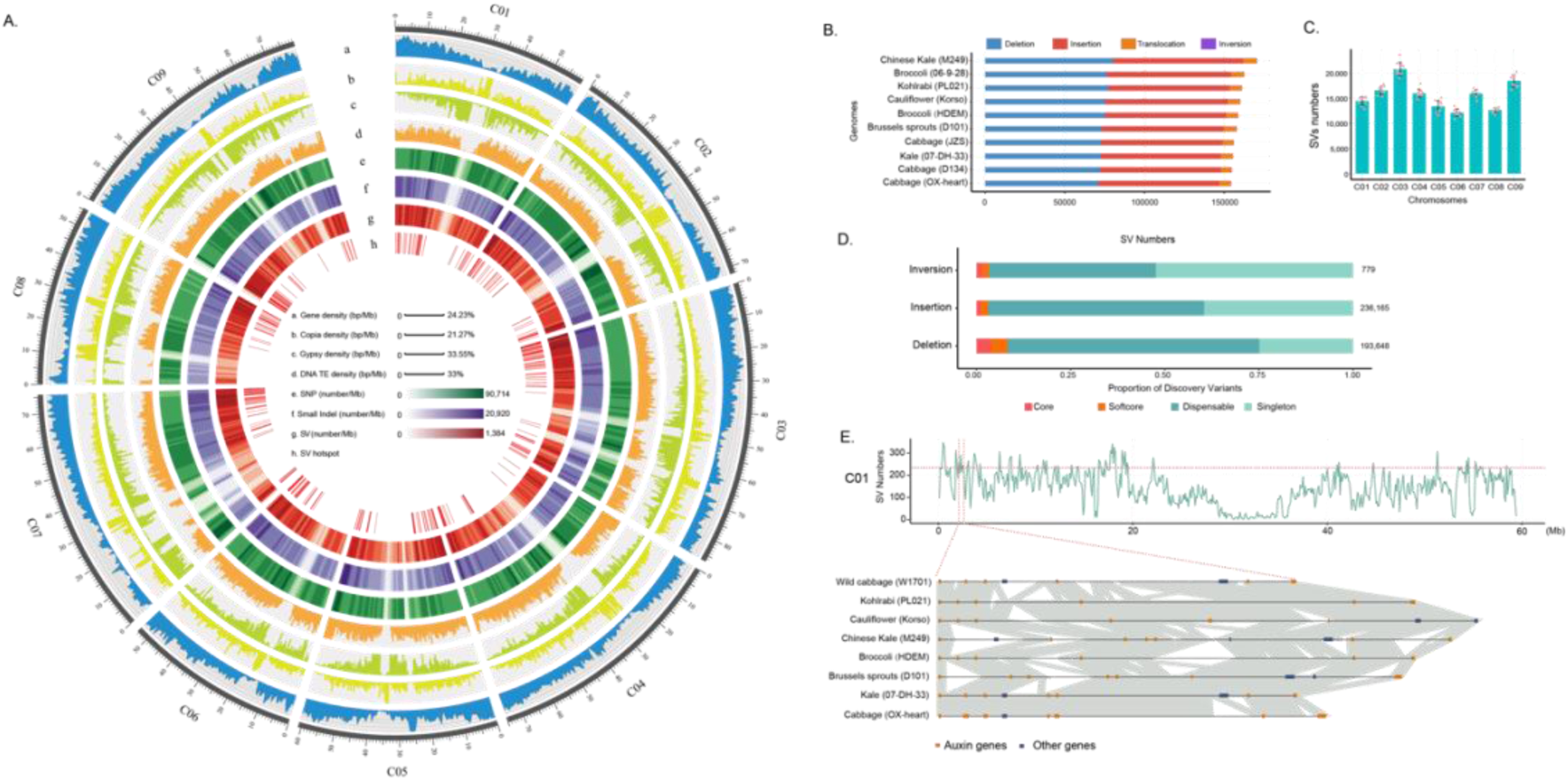
Genome variation landscape in the 11 *B. oleracea* genomes. (A) Circos plot of genomic variants including SNPs, small indels and SVs. (B) Number of different SVs in each of the *B. oleracea* accessions relative to the wild cabbage. (C) Number of SVs across different *B. oleracea* chromosomes. (D) Proportion of SVs (inversions, insertions, and deletions) in different categories of the pan-genome. (E) A representative SV hotspot region on chromosome C01 containing clustered small auxin up-regulated RNA (SAUR) genes.

After merging all SVs across accessions, a total of 469,217 non-redundant SVs were cataloged, including 427,206 indels (236,078 insertions and 191,128 deletions), 41,232 translocations, and 779 inversions. The 427,206 indels made up a total of 371.64 Mb sequences with a mean of 184.97 Mb in each accession, which accounted for ∼32.71% of the assembled genome (Supplementary Table 7). Chromosome C03 harbored the highest number of SVs, while chromosomes C06 and C08 had the fewest SVs (Fig. 3C). Approximately 38.37% of the SVs affected the promoter regions and 2.81% affected the coding sequence regions. Furthermore, 8.67% of the SVs were located in the intron regions (Supplementary Fig. 4A). The SV density in repeat regions was significantly higher than that in non-repeat regions (Supplementary Fig. 4B).

Based on the allele frequency of these SVs in the ten genomes of *B. oleracea* cultivars, we classified them into four categories: core (present in all 10 cultivars), softcore (present in 9 cultivars), dispensable (present in 2-8 cultivars), or singleton (present in only one cultivar). The presence of significantly more dispensable and singleton SVs compared to core and softcore SVs suggests that the *B. oleracea* genome is experiencing rapid recent or ongoing evolutionary changes (Fig. 3E).

The SVs were unevenly distributed across chromosomes, with more than 200 regions identified as SV hotspots (Supplementary Fig. 5A and Supplementary Table 14). This uneven distribution and the presence of SV hotspots highlighted the complex nature of genomic variation in *B. oleracea*. Given the significant variation and diversity in developmental timing and organ formation among different morphotypes, we conducted an analysis of flowering time and auxin-related genes located in the SV hotspots. Of the 631 flowering time genes identified in the whole genome, 165 (26.1%) were located in the SV hotspots (Supplementary Table 10). Similarly, of the 295 auxin-related genes, 79 (26.8%) were found in the SV hotspots (Supplementary Table 11). This provides a genomic basis for the phenotypic differentiation and artificial selection of *B. oleracea*, potentially contributing to the wide range of morphological forms observed in this species. Notably, three SV hotspots on chromosome C01 (2.1-2.6 Mb) (Fig. 3E), C02 (5.2-5.4 Mb) (Supplementary Fig. 5B), and C03 (80.0-80.2 Mb) (Supplementary Fig. 5C), harbored 608, 240, and 256 SVs, respectively, associated with six, five, and five small auxin up-regulated RNA (SAUR) genes, which are early auxin-responsive genes involved in a variety of developmental process, including cell elongation, division, differentiation, and organogenesis. These SV hotspots provide important data foundations for dissecting the link between genomic variations and phenotypic differentiation and organ formation in *B. oleracea*.

### Population structure and domestication of *B. oleracea*

The graph-based pan-genome of *B. oleracea* serves as a foundation for comprehensively investigating population structure, domestication, and molecular mechanisms underlying phenotypic differentiation in *B. oleracea*. Using this graph pan-genome as the reference, we genotyped the 427,206 SVs in 392 *B. oleracea* accessions, including 52 broccoli, 39 Brussels sprouts, 69 cabbage, 116 cauliflower, 37 Chinese kale, 23 kale, 54 kohlrabi accessions, using the genome resequencing data, of which 180 were generated in this study and 212 were from our previous study (Guo et al., 2021) (Supplementary Table 12).

To uncover the domestication history and population structure of *B. oleracea*, we constructed phylogenetic trees and conducted principal component analysis (PCA) using 427,206 SVs and 6,633,641 SNPs, respectively, of the 392 accessions. Both SV and SNP phylogenetic trees, which were rooted with wild cabbage, distinctly divided the accessions into two main clades (Fig. 4A and 4B). Clade I was composed of kale, Brussels sprouts, and cabbage, with kale demonstrating the closest relationship to wild cabbage. This clade exhibited signs of domestication that led to an increase in leafy organs (such as leafy heads). We designated this clade as the “leaf-selected clade”. The second clade encompassed kohlrabi, Chinese kale, broccoli, and cauliflower, with kohlrabi showing the highest genetic similarity to wild cabbage. Morphotypes in this clade displayed a greater degree of phenotypic variation and diverse product organs. Notably, all members of this clade exhibited enlarged stem tissues, such as enlarged tuberous stems in kohlrabi, thickened flower stalks in Chinese kale, and expanded fleshy inflorescence stems in cauliflower and broccoli. This suggests a selection pressure favoring stem development. We referred to this clade as the “stem-selected clade”.

**Fig. 4.**
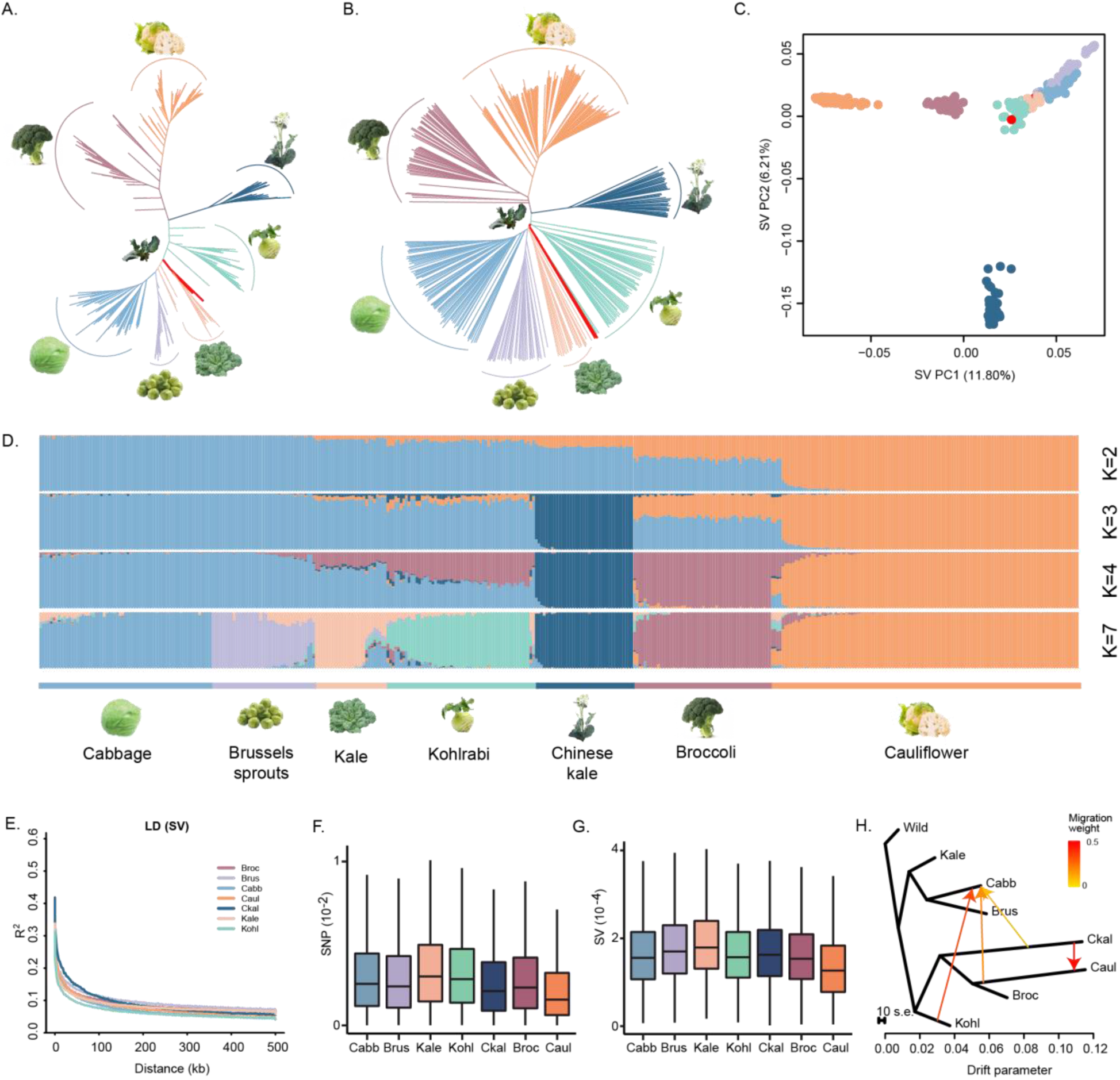
Phylogeny, population structure and gene flow of *B. oleracea*. (A, B) Phylogenetic trees of 392 *B. oleracea* accessions constructed using SNPs (A) and SVs (B). Branch colors correspond to different groups: red, wild cabbage; light orange, kale; purple, Brussels sprouts; light blue, cabbage; cyan, kohlrabi; dark blue, Chinese kale; orange, cauliflower; claret, broccoli. (C) Principal component analysis of *B. oleracea* accessions using SVs. (D) Model-based clustering of the 392 *B. oleracea* accessions with K = 2, 3, 4, and 7. (E) LD decay patterns of different *B. oleracea* morphotypes based on SVs. (F,G) Nucleotide diversity (π) in different *B. oleracea* morphotypes inferred using SNPs (F) and SVs (G). (H) Gene flows among different *B. oleracea* groups inferred by TreeMix. Arrows indicate the direction of migrations. Broc, broccoli; Brus, Brussels sprouts; Cabb, cabbage; Caul, cauliflower; Ckal, Chinese kale; Kohl, kohlrabi.

PCA analysis demonstrated that two wild cabbage accessions were mixed within the kohlrabi and kale groups, further supporting that kohlrabi and kale were most closely related to wild cabbage. The leaf-selected clade morphotypes clustered together. On the other hand, morphotypes in the stem-selected clade exhibited greater genetic distances, consistent with their greater degree of phenotypic variation. In the second principal component, Chinese kale was separated from other cultivated morphotypes, suggesting its relatively independent domestication history (Fig. 4C; Supplementary Fig. 6). This is consistent with the known history of Chinese kale’s domestication in southern China, while other cultivated varieties have origins along the Mediterranean coast (Mabry et al., 2021).

Next, we investigated the genetic structure of the *B. oleracea* population based on the pan-genome SVs and SNPs among the 392 accessions. Under the optimal number of clusters (K=7), all seven different morphotypes could be clearly differentiated (Fig. 4D and Supplementary Fig. 7). The findings from the population structure analysis aligned with those from the phylogenetic and PCA analyses.

We then performed population differentiation analyses using SNP and SV data to calculate *F*_ST_ values for pairwise comparisons of the seven morphotypes. The results demonstrated a consistent trend in *F*_ST_ values between SV and SNP analyses. The smallest *F*_ST_ value was observed between kale and kohlrabi. Despite their distinct clades and phenotypic differences, both represented early domesticated types with a close genetic relationship to wild cabbage. The highest *F*_ST_ value was observed between Brussels sprouts and cauliflower (Supplementary Fig. 8A). Further analyses comparing *F*_ST_ values within and between leaf- and stem-selected clades revealed significant patterns. *F*_ST_ values within the leaf-selected clade were notably lower than those within the stem-selected clade and between the two clades, suggesting less genetic differentiation among leaf-selected morphotypes (Supplementary Fig. 8B and 8C). This highlights the genetic divergence from early domesticated morphotypes like kale and kohlrabi to later-stage variants like Brussels sprouts and cauliflower.

We further analyzed linkage disequilibrium (LD) decay patterns across different cultivated varieties. Kohlrabi displayed the fastest LD decay (Fig. 4E and Supplementary Fig. 9A), consistent with its relatively primitive domesticated type. Kale, representing the relatively primitive group within the leaf-selected clade, showed the highest nucleotide diversity (π), followed by kohlrabi, while cauliflower exhibited the lowest π (Supplementary Fig. 9B and 9C), indicating that cauliflower has undergone more extensive selective breeding. Notably, the trend of π values was similar between SNP and SV analyses across different morphotypes (Fig. 4F and 4G), reinforcing the reliability of these measures of genetic diversity.

### Gene flow among cultivated varieties of *B. oleracea*

We identified four instances of gene flow among the seven cultivar groups of *B. oleracea*: from Chinese kale to cauliflower and broccoli; from kohlrabi, cauliflower and Chinese kale, respectively, to cabbage and Brussels sprouts (Fig. 4H and Supplementary Fig. 10). The gene flow from Chinese kale to cauliflower and broccoli suggests that early developmental transition genes in cauliflower and broccoli may have been derived from Chinese kale. We found 48 flowering-related genes among those transferred from Chinese kale to cauliflower and broccoli. Moreover, we identified auxin-related genes, 14 from Chinese kale, 27 from kohlrabi, and 19 from cauliflower, that were transferred to cabbage and Brussels sprouts (Supplementary Table 13). These genes likely played a role in regulating organ morphogenesis specific to cabbage. Interestingly, despite Chinese kale’s independent evolution in China, separated from Mediterranean-originated cultivars, we found evidence of gene flow from Chinese kale to cauliflower and cabbage, likely resulted from human agricultural activities for variety improvement.

### Population differentiation and leaf head formation

Our pan-genomic and population genetic analyses suggest an early domestication phase in *B. oleracea*, where certain genes and genetic regions were subject to selective pressure. Through SV and SNP *F*_ST_ analyses between the leaf- and the stem-selected groups (Supplementary Fig. 11), we identified 2,815 and 3,462 genes, respectively, under selection, of which 1,654 were identified by both SV and SNP analyses. GO enrichment analysis showed that these selected genes were significantly enriched in biological processes such as stress response, signal transduction, hormone response (particularly auxin), regulation of biological processes, and cell cycle (Supplementary Fig. 12). These results suggested that the initial divergence between leaf and stem organs in *B. oleracea* may have been influenced by specific environmental stress factors, with variations in auxin response likely playing a pivotal role.

Furthermore, we conducted π and XPCLR (Chen et al., 2010) analyses based on SNPs, as well as π and V_ST_ analyses based on SVs, in both leaf- and stem-selected groups. We identified 736 and 645 genes that were specifically selected in the leaf- and stem-selected groups, respectively (Supplementary Table 14). These genes are believed to have played important roles in the early divergence of the two distinct branches of *B. oleracea*. Notably, we discovered a *SAUR8* homolog gene (*C01p004370.1_BolYGL*) that was significantly selected in the leaf-selected group (Fig. 5A). We detected an SV (3267-bp indel) in the promoter region of this gene, with the insertion genotype being predominantly observed in the leaf-selected group while the deletion genotype in the stem-selected group (Fig. 5B). This gene exhibited the highest expression in kale leaves, and its expression was also higher in cabbage leafy heads and Brussels sprouts leaves (Fig. 5C). This insertion was caused by three DNA transposons. Motif analysis of the inserted sequence revealed the presence of multiple transcription activation elements, and light or hormone responses-related motifs (Fig. 5D). These motifs likely contribute to the upregulation of the SAUR gene expression in leaves of kale, cabbage and Brussels sprouts, compared to kohlrabi, Chinese kale, cauliflower, and broccoli. *SAUR* genes are known to play an important role in plant growth and development, particularly in response to the plant hormone auxin (Ren and Gray, 2015; Stortenbeker and Bemer, 2019). The identification of this gene further underscores the potential role of auxin response in the divergence of the leaf- and stem-selected groups.

**Fig. 5.**
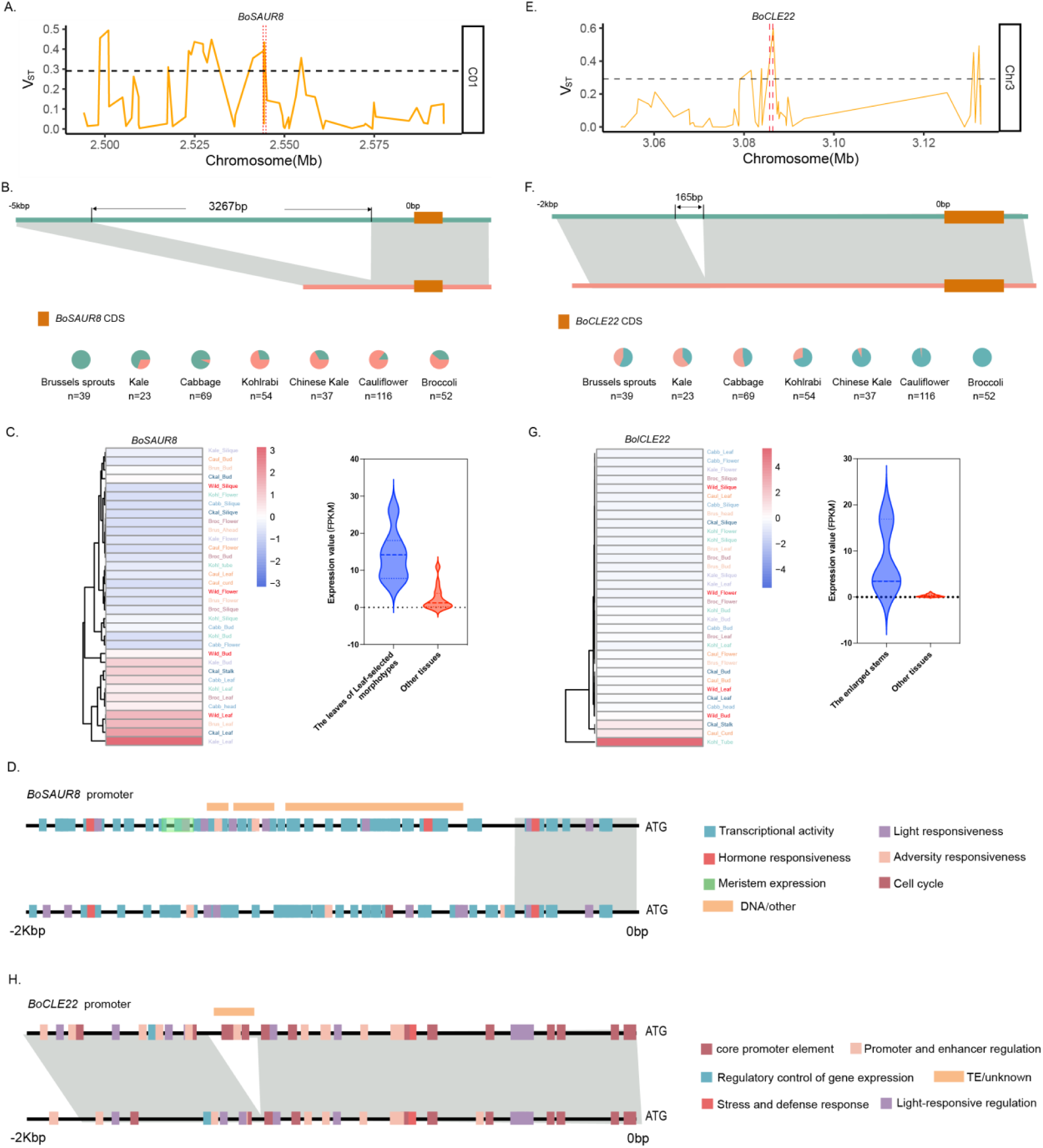
Structural variations of *BoSAUR8* and *BoCLE22*. (A) Selection of *BoSAUR8* in leaf-selected morphotypes inferred by the V_ST_ analysis. (B) An SV (3267-bp indel) in the promoter of *BoSAUR8*. Pie charts show the frequencies of the two alleles of the SV in different morphotype populations. Green: insertion allele; Pink, deletion allele. (C) Expression of *BoSAUR8* in different tissues of different morphotypes (Left), and in leaves and other tissues of leaf-selected morphotypes (Right). (D) Motifs predicted in the promoter region of *BoSAUR8* with different SV alleles. (E) Selection of *BoCLE22* in stem-selected morphotypes. (F) An SV (165-bp indel) in the promoter of *BoCLE22*. Pie charts show the frequencies of the two alleles of the SV in different morphotype populations. Green: insertion allele; Pink, deletion allele. (G) Expression of *BoCLE22* in different tissues of different morphotypes (Left), and in the enlarged stems and other tissues (Right). (H) Motifs predicted in the promoter region of *BoCLE22* with different SV alleles.

We identified a gene from the CLAVATA3/ESR-RELATED (CLE) family, *BolCLE22* (*C03p006260.1_BolYGL*), that showed selective prominence in the stem-selected clade (Fig. 5E). An SV (165-bp indel) in its promoter region displayed clear differentiation across morphotypes, with the insertion genotype predominant in stem-selected varieties (Fig. 5F). *BoCLE22* was prominently expressed in the tuberous stem of kohlrabi, the flower stalk of Chinese kale and the curd of cauliflower, all of which are enlarged stem/peduncle tissues (Fig. 5G). Further analysis revealed that the insertion in the *BoCLE22* promoter was primarily caused by a TE transposition, which introduced motifs related to the core promoter element and promoter and enhancer regulation (Fig. 5H). The *CLE* gene family plays a crucial role in plant growth, especially in orchestrating stem cell homeostasis in shoot apical meristems and vascular cambium meristems (Greb and Lohmann, 2016; Jun et al., 2010; Kang et al., 2022). Our findings underscore the importance of *BoCLE22* in directing meristem development, particularly its influence on early differentiation in stem-centric morphotypes.

In the leaf-selected clade, the transformation from kale to cabbage and Brussels sprouts during domestication is marked by the leaves curling inward to form a leafy head. This change is largely attributed to the interaction between the upper (adaxial) and lower (abaxial) regions during leaf primordium development (Cheng et al., 2016). KAN, a transcription factor from the GARP family, plays a pivotal role in determining abaxial identity and promoting leaf growth (Huang et al., 2014). Our pan-genomic analysis identified four SVs among kale, cabbage, and Brussels sprouts that were associated with the *BoKAN1* gene (Supplementary Fig. 13A), including a notable one in the promoter of *BoKAN1* that showed significant differences between the loose-leaved (kale) and leafy-head (cabbage and Brussels) varieties (Supplementary Fig. 13B). V_ST_ analysis using SVs placed *BoKAN1* in a significantly selected genome region (Supplementary Fig. 13C). Haplotype patterns of these four SVs in kale differed significantly from those in cabbage and Brussels sprouts (Supplementary Fig. 13D). SNP haplotypes of *BoKAN1* also displayed marked differences between loosened leaves and the leafy-head varieties (Supplementary Fig. 13E and 13F). These findings underscore the pivotal role of *BoKAN1* in the evolution of leafy head during the transition from kale to cabbage and Brussels sprouts.

### Evolution of *FLC* genes during *B. oleracea* domestication and improvement

Flowering time is of central importance to determining crop cultivation and harvest seasons to optimize yield and quality. Presence/absence variations of flowering genes increase genetic diversity for adaptation to a wide range of climatic zones and latitudes (Golicz et al., 2016; Schiessl et al., 2018). In *Brassica* crops, the *FLOWERING LOCUS C* (*FLC*) gene is considered a key regulator of vernalization and flowering time (Schranz et al., 2002). The *FLC* gene family in *B. oleracea* consists of four paralogs: *FLC1*, *FLC2*, *FLC3*, and *FLC5*. Our pan-genomic analysis identified a large deletion (9,428 bp) in the genome of Chinese kale, which resulted in the absence of the entire *BoFLC2* gene (Fig. 6A). Genome region harboring this SV showed a strong selection signal between Chinese kale and other morphotypes (Fig. 6B and Supplementary Fig. 14A). Additionally, a single-nucleotide deletion (absence of cytosine) was identified in the fourth exon of *BoFLC2* specifically in cauliflower and broccoli, which underwent purifying selection in these two morphotypes (Supplementary Fig. 14B), presenting in over 90% of the individuals (Fig. 6C). Furthermore, two insertions were identified in the promoter of *BoFLC2* specifically in cauliflower and broccoli, one of which was mediated by a SINE-type transposon (Supplementary Fig. 14C).

**Fig. 6.**
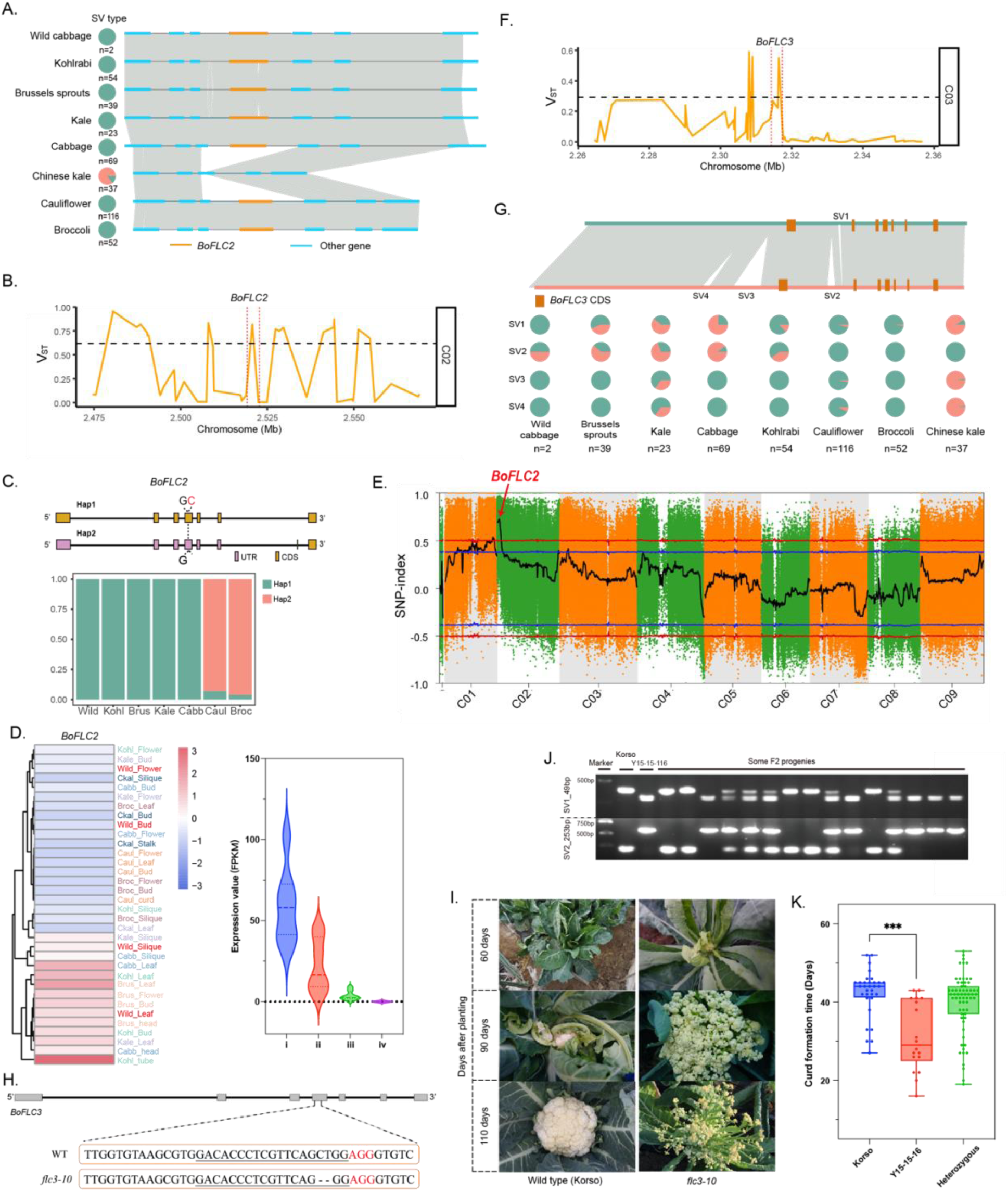
Structural variations of *BoFLC2* and *BoFLC3*. (A) An SV related to *BoFLC2* and its distribution in different morphotype populations. Green: insertion allele; Pink, deletion allele. (B) Selection against *BoFLC2* in Chinese kale inferred by the V_ST_ analysis. (C) A single nucleotide variation in the fourth exon of *BoFLC2* among *B. oleracea*. Green: ‘GC’ allele; Pink, ‘C’ allele. (D) SNP-index analysis between early and late flowering pools of F_2_ progenies from the cross between cauliflower ‘Korso’ and cabbage ‘OX-heart’. (E) SVs related to *BoFLC3* and their distributions in different morphotype populations. (F) Selection of *BoFLC3* in cauliflower and broccoli. (G) Expression of *BolFLC2* in different tissues of different morphotypes (Left), and in vegetative tissues (i) and reproductive tissues (ii) of leaf-selected morphotypes and Kohlrabi, various tissues of cauliflower and broccoli (iii) and Chinese kale (iv). (H) Target sequence and the editing site for knocking out the *BoFLC3* gene using CRISPR/Cas9. (I) Curd at different development stages of wild type ‘Korso’ and *BoFLC3* knockout line *flc3-10*. (J) Genotyping of the 49-bp and 253-bp SVs in the first intron of *BoFLC* in F_2_ progenies of ‘Korso’ and ‘Y15-15-116’. Only results for part of the F_2_ progenies are shown. (K) Curd formation times in F_2_ progenies with different alleles of SVs in the first intron of *BoFLC*.

*BoFLC2* was highly expressed in the leaves of morphotypes in the leaf-selected clade and in the leaves and tubers of kohlrabi. However, its expression was low in all tissues of cauliflower and broccoli, and as expected undetectable in Chinese kale due to the absence of this gene (Fig. 6D). To further support the role of *BoFLC2* in flowering time regulation, an F_2_ mapping population consisting of 1,200 individuals derived from the cross between cauliflower ‘Korso’ and cabbage ‘OX-heart’ was developed. Bulked-segregant analysis combined with genome sequencing (BSA-seq) through comparing the extreme early and late flowering pools pinpointed the genomic region responsible for flowering time variation to a 3-Mb segment on chromosome C02, which notably harbors *BoFLC2* (Fig. 6E). These findings provided insights into the genomic basis of vernalization and flowering trait variations in different cultivated varieties of *B. oleracea*.

Unlike other *Brassica* crops, cauliflower and broccoli have a big flower head (curd), which is composed of a spirally iterative pattern of primary inflorescence meristems with floral primordia arrested (in cauliflower) or flower bud arrested (in broccoli) during their development. Comparative genomic studies have identified the selective variations in the first intron of *BoFLC3* in cauliflower and cabbage (Guo et al., 2021). In this study, population genetic analysis utilizing a graph-based pan-genome approach revealed that *BoFLC3* is subjected to selective pressure on SVs at the species level (Fig. 6F). In comparison to vernalization-dependent cabbage, kale, Brussels sprouts and kohlrabi, two SVs in the first intron of *BoFLC3*, namely SV1_49bp and SV2_253bp, were significantly selected in cauliflower and broccoli, as well as in the vernalization-independent Chinese kale. However, genotypes of these SVs differed among the varieties. Specifically, cauliflower and broccoli showed the SV1_49bp insertion and the SV2_253bp deletion, while Chinese kale exhibited the SV1_49bp deletion and the SV2_253bp deletion (Fig. 6G). Analysis of SNP haplotypes revealed that leaf-selected varieties showed consistent dominant haplotypes, while haplotypes in stem-selected varieties had a greater diversity (Supplementary Fig. 15A). Phylogenetic and evolutionary studies of these haplotypes indicated that the leaf-selected varieties clustered together, cauliflower and broccoli clustered together, Chinese kale stood out as a separate branch, and kohlrabi displayed a mixed pattern, with some individuals clustering with broccoli and others with leaf-selected varieties (Supplementary Fig. 15). To further elucidate the function of *BoFLC3*, we employed CRISPR-Cas9 to knock out *BoFLC3* in the cauliflower cultivar ‘Korso’ (Fig. 6H). The resulting knockout plants exhibited an earlier transition from vegetative to reproductive growth, earlier curd maturation time (∼70 d compared to ∼100 d in wild type ‘Korso’), and a flat curd type compared to the spherical curd in ‘Korso’ (Fig. 6I). These findings strongly supported that *BoFLC3* played a fundamental role in regulating the reproductive transition and flowering in cauliflower, and in the inflorescence propagation and maintenance, which underlie the ‘flower head’ formation in cauliflower and broccoli.

To examine the role of the two SVs in the first intron of *BoFLC3* in cauliflower, we identified a backcross inbred line, ‘Y15-15-116’, from the somatic hybrid progeny of cauliflower and *B. nigra* (Wang et al., 2016). ‘Y15-15-116’ showed an opposite genotype pattern of these two SVs compared to the recipient parent cauliflower ‘Korso’, the SV1_49bp deletion and the SV2_253bp insertion. Interestingly, the curd formation, maturation, and flowering of ‘Y15-15-116’ were significantly earlier than those of ‘Korso’, but similar to those in the *BoFLC3* knockout plants (Supplementary Fig. 16A and 16B). It’s worth noting that when the floral bud of ‘Y15-15-116’ had already developed into a well-formed flower primordium, ‘Korso’ still remained at the inflorescence primordium stage (Supplementary Fig. 16 C). The expression of *BoFLC3* during the entire curd development period in ‘Y15-15-116’ was significantly lower than that in ‘Korso’ (Supplementary Fig. 16D). To further support the role of these two SVs in curd development, we constructed an F_2_ population containing 130 lines using ‘Korso’ and ‘Y15-15-116’ as the parents. We found that in the F_2_ population, the inflorescence formation and flowering time of individuals with the ‘Korso’ SV genotype (characterized by SV1_49bp insertion and SV2_253bp deletion) were significantly later than those with the ‘Y15-15-116’ SV genotype (characterized by SV1_49bp deletion and SV2_253bp insertion) (Fig. 6J and 6K). Together these results proved that the two SVs in the first intron of *BoFLC3* play an important functional role in the regulation of flowering and curd development.

## Discussion

In this study, we generated high-quality genome assemblies for the wild cabbage and five different morphotypes of *B. oleracea*, including kale, kohlrabi, broccoli, Brussels sprouts, and Chinese kale. Together with five recently released high-quality genomes from broccoli and other two *B. oleracea* morphotypes, cabbage and cauliflower, we constructed a graph-based pan-genome of *B. oleracea* that captured ∼469 K SVs, including ∼427 K indels, ∼41 K translocations and 779 inversions. The newly generated genome assemblies and pan-genome add comprehensive and valuable resources of *B. oleracea* that will facilitate mining nature variations in the germplasm and accelerate future functional studies and molecular breeding of this important crop species with various morphotypes. Our population analyses based on pan-genomics reveal two major distinct lineages of *B. oleracea* morphotypes, the leaf-selected and stem-selected clades, supporting that early domestication of *B. oleracea* had been bifurcated into two main pathways centering on the selection for enhanced leaf and stem features, respectively. Cai et al. (2022) have postulated that cabbage and cauliflower originate from different kale clades, leading to two domestication lineages: the ‘leafy head’ lineage and the ‘arrested inflorescence’ lineage (Cai et al., 2022). Expanding upon this understanding and based on our population genomic analyses leveraging the graph-based pan-genome, we propose that the initial differentiation of *B. oleracea* was influenced by artificial selection of leaves and stems, respectively. The ‘leafy head’ and the ‘arrested inflorescence’ lineages proposed in Cai et al. (2022) (Cai et al., 2022) are derived from further domestications within the leaf- and stem-selected clades, respectively, in terms of leaf morphology and the development of flowering and inflorescence. Furthermore, gene flows were detected from Chinese kale to cauliflower and broccoli, and from stem-selected morphotypes to cabbage. These gene flows are likely the results of human agricultural activities and ongoing variety improvement processes, which have further shaped the genetic diversity and structure of *B. oleracea*.

Our pan-genome and population analyses further reveal the potential molecular mechanisms underlying the initial domestication of *B. oleracea* and presents a comprehensive molecular regulatory framework elucidating the domestication and organogenesis processes in *B. oleracea* (Fig. 7). Our results have pinpointed the potential important roles of SVs in the promoters of *BoSAUR8* and *BoCLE22* in the early stage of *B. oleracea* domestication, which are involved in the auxin-mediated response process and the regulation of stem cell differentiation, respectively. Such molecular changes facilitated the selective differentiation of leaves and stems, which had subsequently led to the domestications that gave rise to kale and kohlrabi, respectively. Within the clade characterized by leaf selection, the SVs in the promoter of the *BoKAN1* gene, a crucial determinant of leaf abaxial identity, could play an important role in inducing leaf inward curling. These genetic alterations further pave the way for the domestication toward cabbage and Brussels sprouts, both of which exhibit terminal and axillary buds that culminate in the formation of a leafy head. In the stem-selected clade, mutations in the *FLC* genes were found to be involved in regulating vernalization and flowering times. Such genetic variations have resulted in the emergence of varieties characterized by floral/inflorescence stems. A notable instance is the absence of the *BoFLC2* gene in Chinese kale, which could have resulted in its early flowering and independent of vernalization, leading to the formation of floral stalks as its primary yield. Furthermore, the deletion of a single base in the fourth exon of *BoFLC2*, coupled with SVs in the first intron of *BoFLC3*, could play critical roles in influencing the precocious reproductive transformation and curd development observed in cauliflower and broccoli. Overall, the pan- and population genomic analyses of different morphotypes shed light on the domestication and differential organogenesis of *B. oleracea*, providing valuable insights into the evolutionary history, genetic variation, and functional aspects of this important vegetable species. Furthermore, results from this study have implications for future breeding and genetic improvement in *B. oleracea* and contribute to our broader understanding of crop domestication processes.

**Fig. 7.**
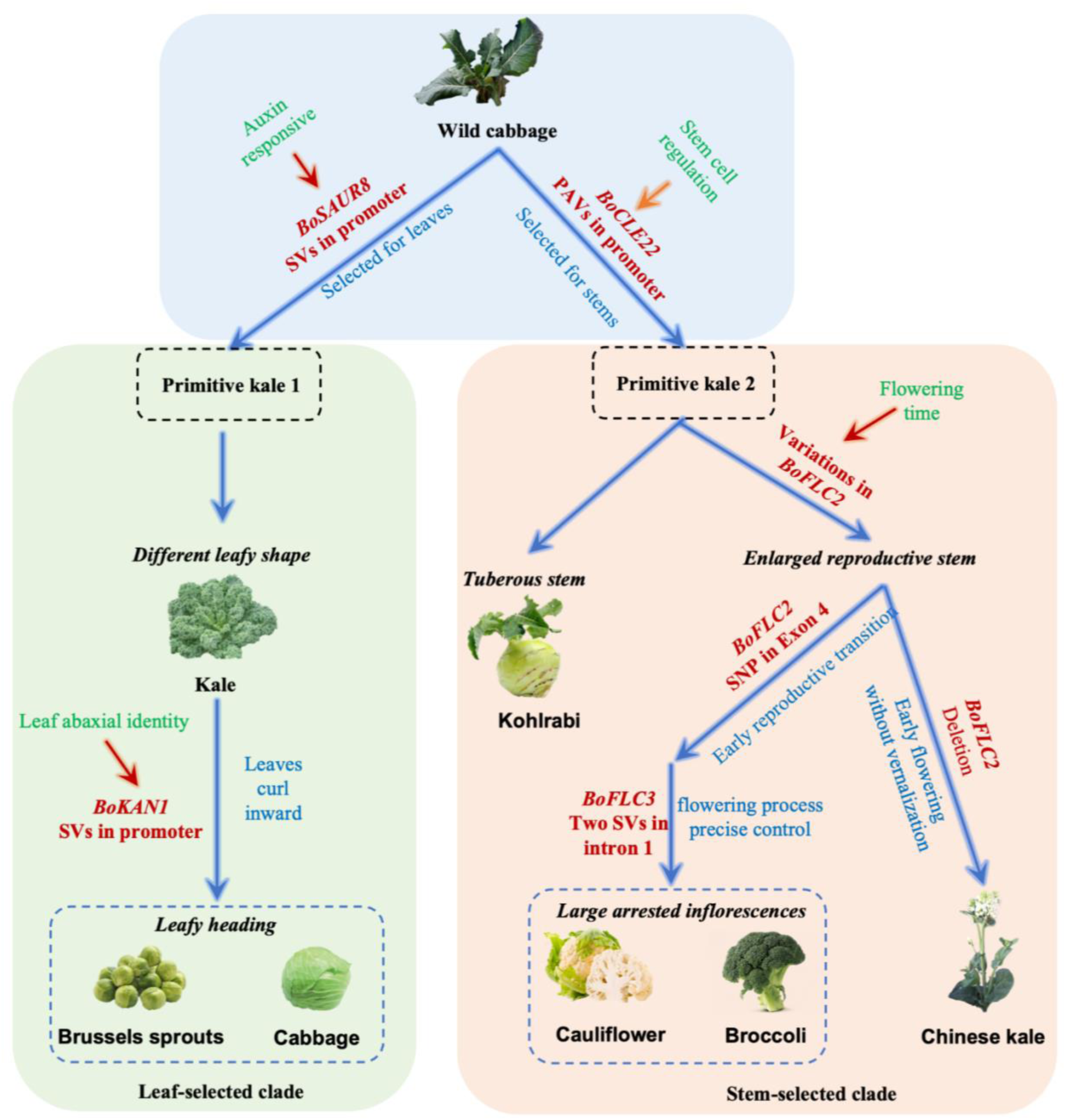
Model of *B. oleracea* domestication and organ development.

## Materials and Methods

### Plant materials

For genome assembly, the wild cabbage accession (*Brassica oleracea* W1701) used in this study is an inbred line from Oil Crops Research Institute of the Chinese Academy of Agricultural Sciences, the Broccoli accession (var. *italica* 06-9-28) is a highly inbred line from Tianjin GengYun Seed Company, the kohlrabi accession (var. *gongylodes* PL021) is a highly inbred line from Vegetable Research Institute of Jiangsu Academy of Agricultural Science, the Brussels sprouts accession (var. *gemmifera* D101) is a highly inbred line from Vegetable Research Institute of Jiangsu Academy of Agricultural Science, the kale accession (var. *acephala* 07-DH-33) is a double haploid (DH) line from Beijing Vegetable Research Center of Beijing Academy of Agriculture and Forestry Science, and the Chinese kale accession (var. *alboglabra* M249) is a highly inbred line from Beijing Vegetable Research Center of Beijing Academy of Agriculture and Forestry Science. All seedlings were planted in the greenhouse of Beijing Vegetable Research Center of Beijing Academy of Agriculture and Forestry Science.

### Genome sequencing and assembly

Young flesh leaves were collected from a single plant for each of the six *B. oleracea* accessions. Genomic DNA was extracted using the QIAGEN Genomic-tips (QIAGEN®). A 15-kb library was constructed for each accession and sequenced on the PaciBio Sequel II platform by NextOmics Biosciences Co., Ltd. (Wuhan, China). HiFi/CCS reads of each accession were generated using the ccs software v.3.0.0 (https://github.com/PacificBiosciences/ccs).

HiFi reads from each of the six accessions were assembled into contigs using hifiasm (v0.12) (Cheng et al., 2021). The assembled contigs were corrected using Illumina short reads with Nextpolish (v1.2.4) (Hu et al., 2020) with default parameters, and then further assembled into pseudochromosomes using RaGOO (v1.1) (Alonge et al., 2019) with the ‘Korso’ genome used as the reference. BUSCO (Simão et al., 2015) and LTR Assembly Index (LAI) (Ou et al., 2018) were used to determine the completeness of the assemblies based on the Embryophyta Plant database and full-length long terminal repeat retrotransposons, respectively.

### Genome annotation

A de novo repeat library was constructed for each genome assembly using LTR_FINDER (Xu and Wang, 2007), RepeatScout (http://www.repeatmasker.org/), and Repeat-Modeler (http://www.repeatmasker.org/RepeatModeler.html), which was combined with the RepBase database (http://www.girinst.org/repbase). The combined repeat library was then used to scan each genome assembly using RepeatMasker (http://repeatmasker.org/) to identify repetitive sequences.

Protein-coding genes were predicted from each of the repeat-masked genomes using a comprehensive strategy combining ab initio, protein homology, and transcriptome-based predictions. For protein homology prediction, protein sequences from *B. napus* (ZS11), *B. juncea* (SCHZ), *B. rapa* (Chiifu), *B. nigra* (YZ12151), *B. oleracea* (Korso and OX-heart), and *Arabidopsis thaliana* were aligned to the corresponding genomes using BLAST (She et al., 2009) with an E-value cutoff of 1e^-5^ and the hits were joined using SOLAR (Yu et al., 2006). GeneWise (Birney et al., 2004) was used to predict the exact gene structures of the corresponding genomic regions having BLAST hits. For ab initio gene prediction, five programs, AUGUSTUS (v2.5.5) (Stanke et al., 2006), GENSCAN (v1.0) (Burge and Karlin, 1997), GENEID (Guigo, 1998), GlimmerHMM (v3.0.1) (Majoros et al., 2004) and SNAP (Korf, 2004), were employed. For transcriptome-based prediction, RNA-seq data were mapped to the assembly using HISAT2 (Kim et al., 2019), and then assembled into transcripts using a reference-guided approach with StringTie (Pertea et al., 2015). RNA-seq data were also *de novo* assembled into transcripts using Trinity (Grabherr et al., 2011). The complete coding sequences (CDS) were predicted from the assembled transcripts using PASA (Haas et al., 2003). Gene models from the above ab initio, protein homology, and transcriptome-based predictions were integrated using EVidenceModeler (EVM) (Haas et al., 2008) to obtain a final set of high-confidence gene models for each genome assembly.

The predicted protein-coding genes were functionally annotated by comparing their protein sequences against the following databases: GenBank nr, the Kyoto Encyclopedia of Genes and Genomes (KEGG), and Swiss-Prot using BLAST with an E-value cut-off of 1×10^−5^. In addition, the protein sequences were also compared against the InterPro domain database using InterProScan (Hunter et al., 2009) with the parameters ‘-f TSV -dp -gotermes -iprlookup’.

### Pan-genomic gene annotation improvement

In order to keep gene predictions of the eleven *B. oleracea* genomes used for pan-genome construction at the same standard, we improved the gene predictions of these genomes using a unified pipeline. First, we carried out pairwise collinearity analysis for the 11 assemblies using MCScanX (Wang et al., 2012) with default parameters. We used genes from the wild cabbage as the base, and genes from other 10 assemblies were added in a stepwise manner: a gene was added into the gene list and assigned a new locus ID if it had no collinear genes in the gene list generated by the preceding step. This process was repeated until all genes from 10 assemblies were added. All genes resulting from the above process were remapped to the 11 assemblies using GMAP (version 2018-07-04) (Wu and Watanabe, 2005) with parameters ‘--format=gff3_gene -n 10 --min-trimmed-coverage=90 --min-identity=90’. Genes not overlapping with the original predicted genes in the genome were added. We further classified these gene models into high-confidence (HC) and low-confidence (LC) protein-coding genes based on a stringent confidence classification method. Briefly, BLASTP was used to align the predicted protein sequences to protein datasets of *B. oleracea*, *B. napus*, *B. rapa*, *B. nigra*, *B. juncea*, *Oryza sativa* and *A. thaliana* with an e-value cutoff of 1e^-10^. Genes were designated high confidence (HC) if they had significant BLAST hits to at least two reference proteins with a similarity above a threshold (>60% for *O. sativa* and *A. thaliana*; >80% for *B. rapa*, *B. nigra*, *B. juncea*; and >90% for the *B. napus*). The high-confidence (HC) gene set was kept and used for subsequent analyses.

### Construction of the *B. oleracea* gene family-based pan-genome

Predicted protein sequences from the 11 *B. oleracea* genomes were clustered into gene families using OrthoFinder (v1.1.4) (Emms and Kelly, 2015). Gene families containing members from all 11 *B. oleracea* accessions were defined as core gene families, those containing members from ten accessions were defined as softcore gene families, those with members from 2 to 9 *B. oleracea* accessions were defined as dispensable gene families, and those that were present in only one accession were defined as private/specific gene families.

### Genomic structural variation detection

The 10 *B. oleracea* genomes were aligned to the wild cabbage genome using NUCmer (v3.23) (Kurtz et al., 2004) with parameters ‘--mum --maxgap=500 --mincluster=1000’. The alignment results were used as input for three SV detection programs, Assemblytics (Nattestad and Schatz, 2016), SyRI (Goel et al., 2019) and SVMU (Chakraborty et al., 2019), all of which were run with default parameters. The resulting three raw SV sets for each accession were merged using the strategy similar to that described in a previous study (Qin et al., 2021). Briefly, two SVs were merged if the overlapping ratio of these two SVs (the length of the overlapping sequence/the length of non-redundant genome segment covered by the two SVs) exceeded 90%. For inversions and translocations, we only considered candidates called by SyRI. We then merged SVs (including indels, inversions and translocations) from the 10 accessions relative to the wild cabbage. For the deletion SVs, if the overlapping ratio of the two SVs exceeded 90%, then these two SVs were merged into one. For insertion SVs from the 10 accessions, if their overlapping ratio (the length of the shorter SV/the length of longer SV) exceeded 90%, and the difference of their positions on the wild cabbage genome was less than 5 bp, the two SVs were merged into one. The merged non-redundant SVs and wild cabbage genome were used to construct graph-based pan-genome using the Variation graph toolkit (Garrison et al., 2018).

### Genome re-sequencing, read mapping and SNP calling

A total of 180 *B. oleracea* accessions (Supplementary Table 12) were self-pollinated over multiple generations before re-sequencing. Genomic DNA extracted from fresh leaves was used for the construction of Illumina libraries, which were sequenced on the Illumina HiSeq X Ten platform. A total of 2.9 Tb (∼16.14 Gb per sample) of cleaned data was generated after processing the raw reads with the fastp software (Chen et al., 2018) with default parameters. The paired-end reads of the 180 accessions from this study and 212 accessions from our previous study (Guo et al., 2021) were mapped to the wild cabbage genome using BWA (v0.7.8) (Li and Durbin, 2009) with the command ‘mem --t 4 --k 32 --M’. Duplicated reads were removed with SAMtools (v.0.1.19) (Li et al., 2009). Genomic variants were then identified with the HaplotypeCaller and the GenotypeGVCFs functions in Genome Analysis Toolkit (GATK) (McKenna et al., 2010). Raw SNPs were filtered with the following parameters: depth ≥ 4, genotype quality ≥ 5, minor allele frequency (MAF) ≥ 0.05 and missing rate ≤ 0.1. The identified SNPs were further annotated using ANNOVAR (v2013-05-20) (Wang et al., 2010).

### SV genotyping

We genotyped 469,217 SVs using the short sequencing reads of the 392 *B. oleracea* accessions using the SV genotyping pipeline integrated in the Variation graph toolkit (Garrison et al., 2018) through the command of ‘map’, ‘pack’ and ‘call’ with default parameters, with the graph-based pan-genome as the reference. BCFtools (Danecek et al., 2021) was used to merge SVs of all accessions, and then VCFtools (Danecek et al., 2011) was used to filter SVs with the parameters ‘--mac 5 --minDP 4 --minQ 100 --max-missing 0.8’.

### Population structure and phylogenetic analyses

The population structure of *B. oleracea* was determined using the program ADMIXTURE (v1.23) (Alexander et al., 2009) with K values (the putative number of populations) from 2 to 10. The K value of 7 was chosen because clusters maximized the marginal likelihood. To construct maximum-likelihood phylogeny, we used 652,440 independent SNPs obtained using PLINK (Purcell et al., 2007) with parameters ‘--const-fid --indep-pairwise 50 5 0.26’. Phylogenetic tree was constructed using IQ-TREE (v 2.0.3) (Nguyen et al., 2015), based on the best model determined by the Bayesian information criterion. Bootstrap support values were calculated using the ultrafast bootstrap approach (UFboot) with 1,000 replicates. The phylogenetic tree was visualized using the online tool EvolView (https://www.evolgenius.info/evolview/). PCA was performed using GCTA (Yang et al., 2011). Nucleotide diversity (π) and population fixation index (*F*_ST_) were calculated using VCFtools (Danecek et al., 2011). To estimate and compare the pattern of LD decay among different groups, the squared correlation coefficients (*r*^2^) between pairwise SNPs were computed using the PopLDdecay (v.3.40) (Zhang et al., 2019) software with the parameters ‘--MaxDist 500 --MAF 0.05 --Miss 0.1’. All the analyses were also performed using SVs.

### Introgression and demographic history

The population relatedness and migration events were inferred using TreeMix (Pickrell and Pritchard, 2012). We constructed the tree with the wild cabbage as the root, and then ran TreeMix with the number of migration events from 1 to 6. To detect admixture, we computed D-statistics (Durand et al., 2011) based on ABBA and BABA SNP frequency differences. For a triplet of taxa P1, P2 and P3, and an outgroup O, that follows the phylogeny of (((P1, P2), P3), O), a D statistic significantly different from zero indicates P3 exchanged gene with P1 (D value <0) or P2 (D value >0). The f-branch statistic was then used to infer introgressions among the subgroups using the software package Dsuite (Malinsky et al., 2021). The *fd* statistic (Martin et al., 2015) was used to calculate the fraction of introgression, which signifies gene flow when 0 < fd < 1.

### Selective-sweep analysis

The XP-CLR score were calculated based on SNPs using the XP-CLR package (Chen et al., 2010) with sliding windows of 10 kb and a step size of 1 kb. Genome regions with top 5% XP-CLR scores were identified as the candidate selective sweeps, and genes in these regions were considered as candidate genes under selection.

We calculated V_ST_ to identify divergent SV profiles between populations. V_ST_ was calculated as follows:

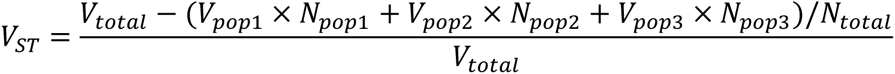

where V_total_ is the total variance, V_pop_ is the SV variance for each respective population, N_pop_ is the sample size for each respective population, and N_total_ is the total sample size. SVs with the top 5% V_ST_ values were identified as candidate selective SVs, and genes with distance less than 5 kb to the selective SVs were considered as candidate genes under selection.

Genome regions with top 5% of nucleotide diversity (π) or fixation index (*F*_ST_) were also considered as candidate selective regions and genes in these regions were considered as candidate genes under selection.

### Transcriptome sequencing and data processing

Tissues of wild cabbage (leaf, bud, flower, and silique), cauliflower (leaf, curd, bud, and flower), broccoli (leaf, curd, bud, and flower), cabbage (leaf, leafy head, flower, bud, and silique), Brussels sprouts (leaf, axillary leafy head, bud, and flower), Chinese kale (leaf, flower stalk, bud, and silique), kale (leaf, bud, flower, and silique), and kohlrabi (leaf, tuberous stem, bud, flower, and silique) were collected for transcriptome sequencing. Three biological replicates were conducted for each sample. Leaves were sampled at the rosette stage of plants with 8-10 leaves (4-6 weeks after planting). The leafy heads, curds, tuberous stems, and flower stalk were sampled at the maturity period of these reproductive organs. The buds were about 2 mm in length, the flowers were blooming, and the siliques were at the developing stage including seeds. RNA was extracted from each tissue using the TIANGEN RNAprep Pure Kit (Cat: No. DP441). RNA quality was assessed using an Agilent 2100 BioAnalyzer. RNA-Seq libraries were prepared using the Illumina TruSeq RNA Sample Prep Kit and sequenced on an Illumina HiSeq X Ten system. Raw RNA-Seq data were preprocessed using the NGS QC Toolkit (v2.3.3) (Patel and Jain, 2012) to remove adaptors, low-quality bases, and reads containing more than 10% unknown bases (“N”). The cleaned reads were mapped to the reference genomes using HISAT2 (Kim et al., 2019) with default parameters. Based on the alignments gene expression levels were derived and represented as fragments per kilobase of transcript per million mapped fragments (FPKM).

### BSA-seq analysis

An F_2_ mapping population consisting of 1,200 individuals derived from the cross between cauliflower ‘Korso’ and cabbage ‘OX-heart’ was constructed. In the Fall of 2021, the entire F_2_ population was planted in the greenhouse at Beijing Vegetable Research Center of Beijing Academy of Agriculture and Forestry Science, and an investigation of the flowering time of the individuals was conducted. Twenty individuals with the earliest flowering time and twenty with the latest flowering time were selected. DNA was extracted from young flesh leaves of these plants, and equal amounts were mixed to create early- and late-flowering DNA bulks. High-throughput resequencing was performed on the Illumina platform, resulting in 14.2 Gb and 14.8 Gb of cleaned data for early- and late-flowering bulks, respectively. The sequencing reads were aligned to the genome of ’Korso’ using BWA-MEM (Li et al., 2009). Variant calling was carried out using GATK UnifiedGenotyper (v3.5) (DePristo et al., 2011). SNPs and small indels were filtered using GATK VariantFiltration function with parameters: -Window 4, -filter ‘QD < 4.0 || FS > 60.0 || MQ < 40.0’, -G_filter ‘GQ < 20’. The SNP-index and the Δ(SNP-index) calculation and QTL identification were performed according to the methods described in Takagi et al. (2013).

### *BoFLC3* knockout by CRISPR/Cas9 in cauliflower ‘Korso’

The *BobFLC3* genomic sequence of ‘Korso’ was utilized to design sgRNAs using the online tool CRISPR-P 2.0 (Liu et al., 2017). The sequences of sgRNAs are listed in Supplementary Table 15. The vector construction was conducted in accordance with the methodology outlined in Xing et al. (2014) (Xing et al., 2014). The accurately sequenced vector was then introduced into *Agrobacterium* and stored at -80 °C. The cauliflower transformation was performed using the protocol established by Wang et al. (2022) (Wang et al., 2022a). Transgenic plants were evaluated for their resistance to glufosinate using the BASTA (glufosinate) resistance test strips. The editing types in the transgenic plants were identified through analysis of high-throughput sequences using the online tool Hi-TOM (Liu et al., 2019). The validated edited plants were grown in the greenhouse at Beijing Vegetable Research Center of Beijing Academy of Agriculture and Forestry Science in the Fall of 2021 and 2022, and their characteristics such as curd development and flowering time were investigated.

### Genotyping of SVs in the first intron of *BoFLC3*

In the spring of 2023, the F_2_ population from the cross between ‘Korso’ and ‘Y15-15-116’ were planted in open fields to record the flowering time of all individuals in the population. To genotype the two SVs located in the first intron of *BoFLC3* in this F_2_ population, primers were designed on both sides of each of the two SVs (Supplementary Table 15). Electrophoresis was conducted using a 2% agarose gel, a voltage of 90V, and a duration of 45 minutes.

## Data availability

Raw PacBio, transcription, and re-sequencing reads have been deposited in the National Center for Biotechnology Information (NCBI) BioProject database under the accession number PRJNA1030743. Genome assemblies and annotations of wild cabbage (W1701), broccoli (06-9-28), kohlrabi (PL021), kale (07-DH-33), Brussels sprouts (D101), and Chinese kale (M249) are available at the National Genomics Data Center (NGDC) (https://ngdc.cncb.ac.cn/) under the accession number PRJCA020663. All data will be made public upon the acceptance of this manuscript.

## Funding

This research was supported by grants from the National Key Research and Development Program of China (2022YFF1003001), National Natural Science Foundation of China (32072576), National Modern Agriculture Industry Technology System (CARS-23-G42), Jiangsu Provincial Key Research and Development Program (BE2021376), the Innovation Program of the Beijing Academy of Agricultural and Forestry Sciences (KJCX20230121), and the Collaborative Innovation Program for Leafy and Root Vegetables of the Beijing Vegetable Research Center, Beijing Academy of Agriculture and Forestry Science (XTCX202302).

## Author contributions

F.L., J.L., Z.F. and N.G. conceived and designed the research. F.L. and J.L. managed the project. N.G., L.C., T.W. and C.J. performed data analysis. F.L., J.L., S.W., D.Z. and S.H. collected samples. M.D., M.Z., GW., L.M., X.L., N.G., S.W. and H.X. performed experiments. N.G., F.L. and L.C. wrote the manuscript. Z.F., F.L., J.L., N.G., S.W., L.C. revised the manuscript.

## Acknowledgements

We thank Xiaoming Wu and Heliang Ji for providing seeds of wild cabbage (M1701) and broccoli (06-9-28), Weiming He and Pangyuan Liu for providing seeds of some resequencing accessions, and Feng Cheng for discussion and advice.

## Conflict of interest

The authors declare no conflict of interest.

## Notes

### Competing Interest Statement

The authors have declared no competing interest.

