## Supplementary figures for "Pan-genome analysis of different morphotypes reveals genomic basis of *Brassica oleracea* domestication and differential organogenesis"

**­­­=**
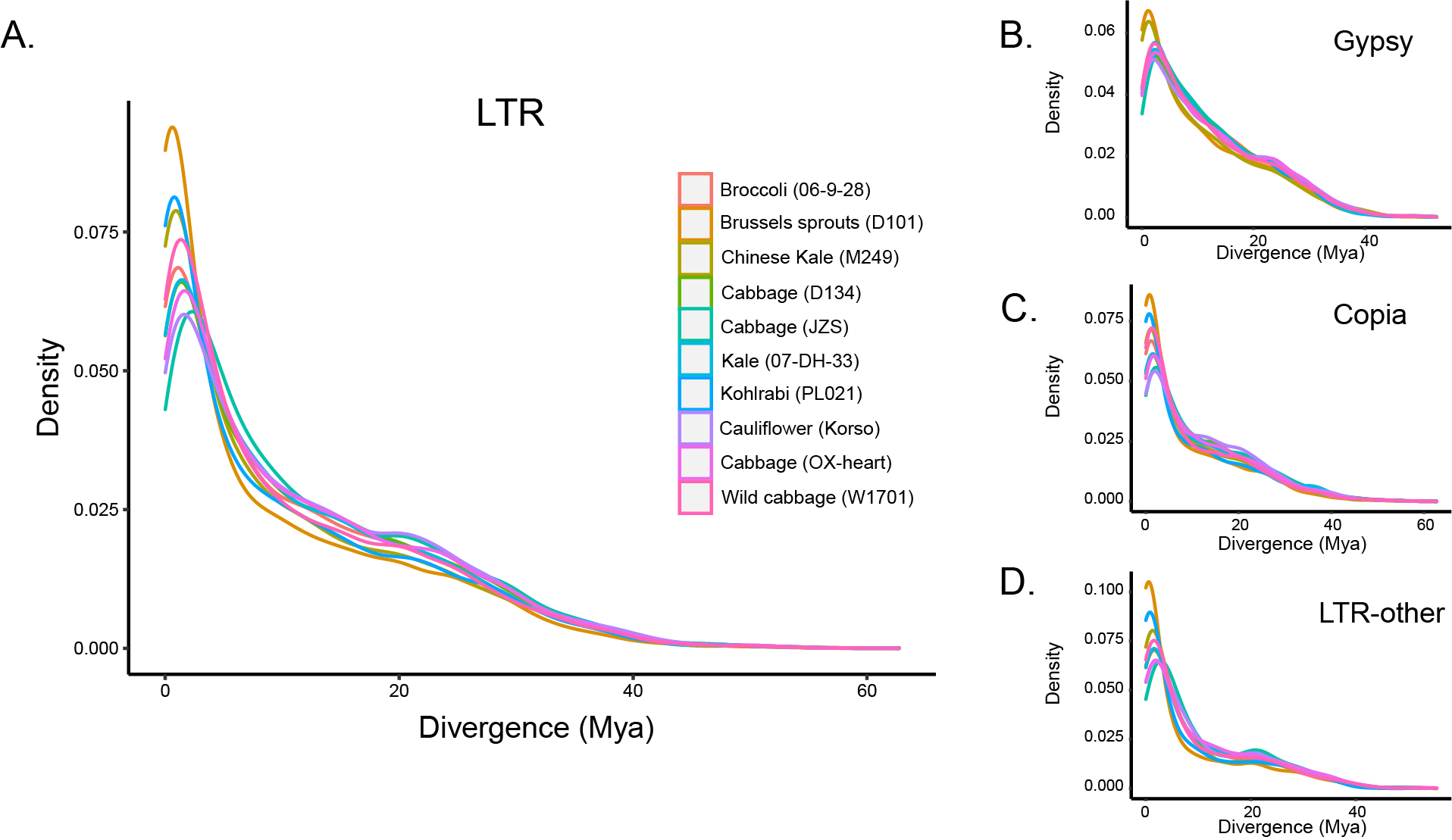

**Supplementary Fig. 1** Estimated insertion times of all intact LTR (A), Gypsy (B), and Copia (C) and other LTR (D) retrotransposons in the 10 *B. oleracea* genomes.

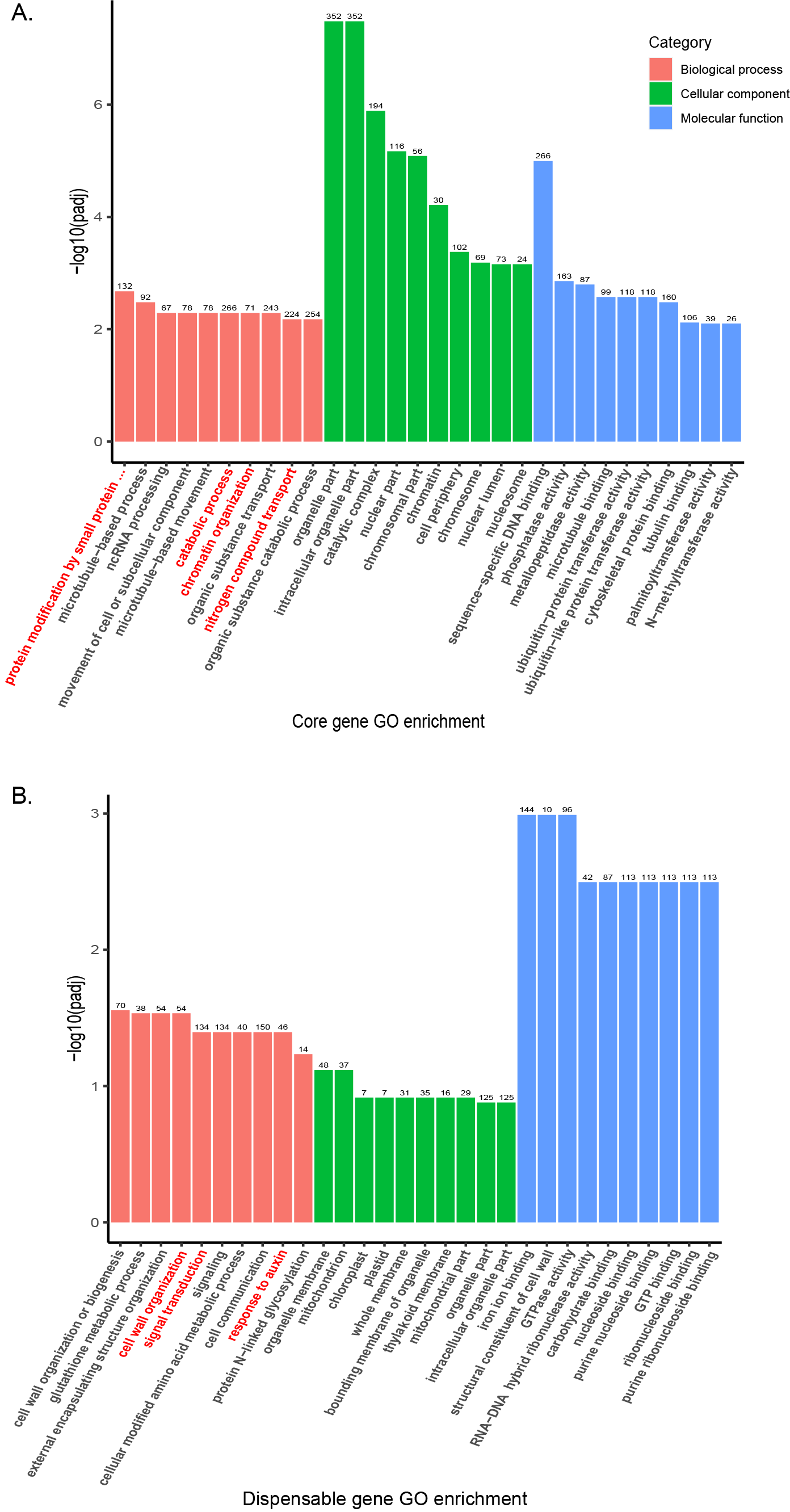

**Supplementary Fig. 2** GO enrichment analyses of core (A) and dispensable (B) genes in the *B. oleracea* pan-genome.

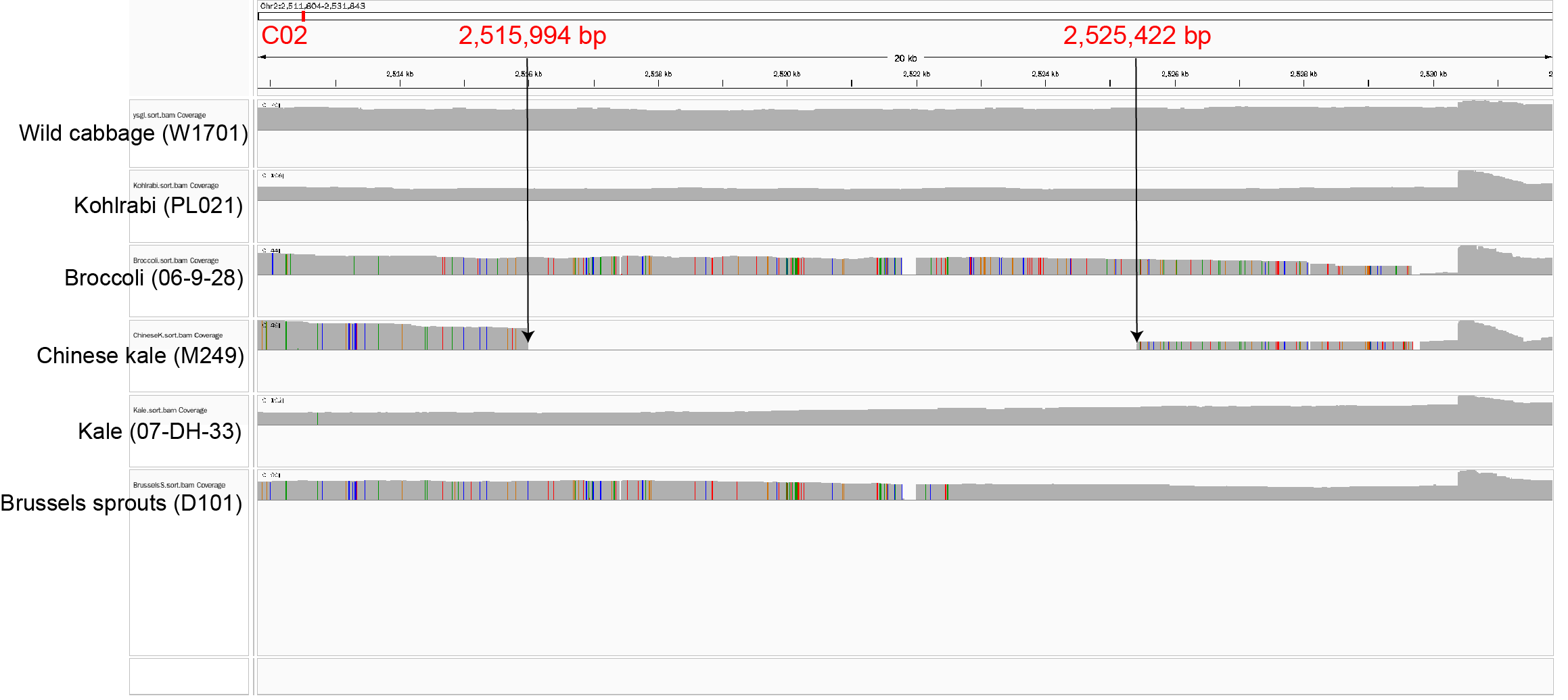

**Supplementary Fig. 3** An example of manual SV validation. A 9,428-bp deletion in Chinese kale was confirmed by visualizing the alignments of PacBio long reads to the wild cabbage assembly.

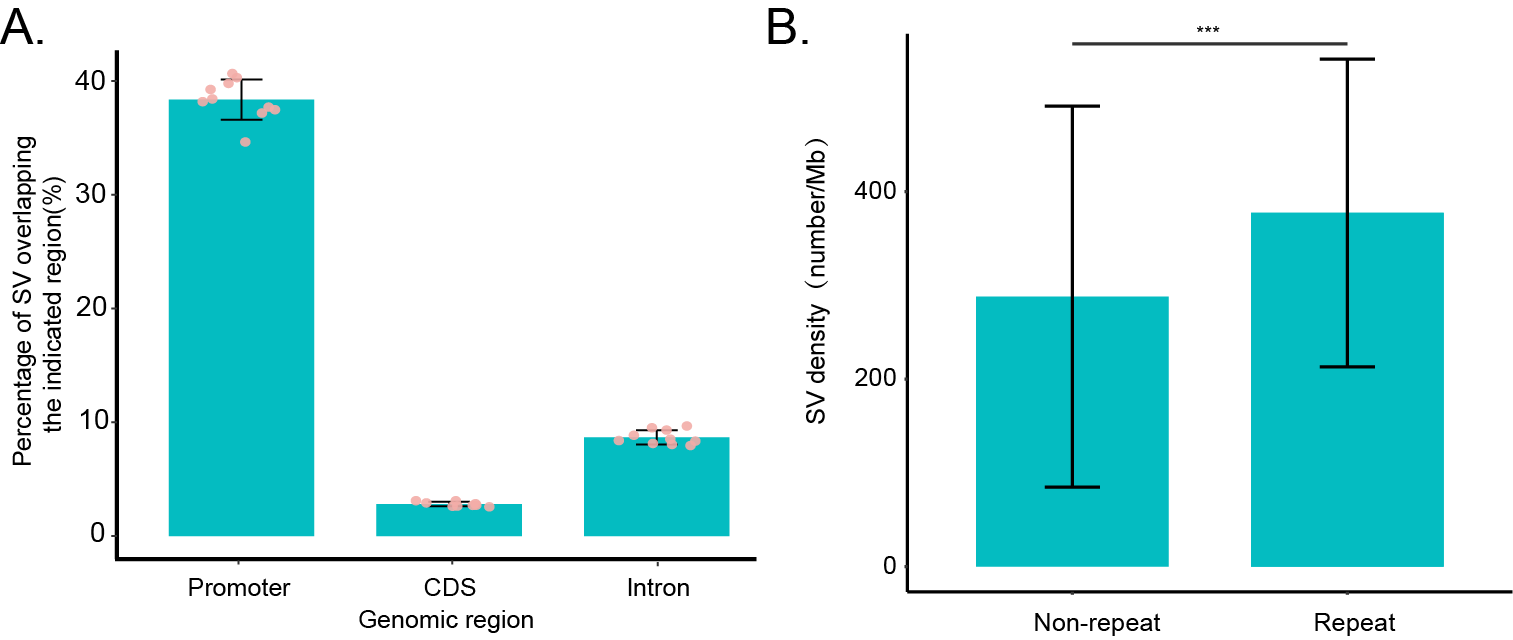

**Supplementary Fig. 4** SVs associated with different genome features. (A) Percentage of SVs overlapping with the promoter, CDS, and intron regions. (B) SV densities in repetitive and non-repetitive genome regions.

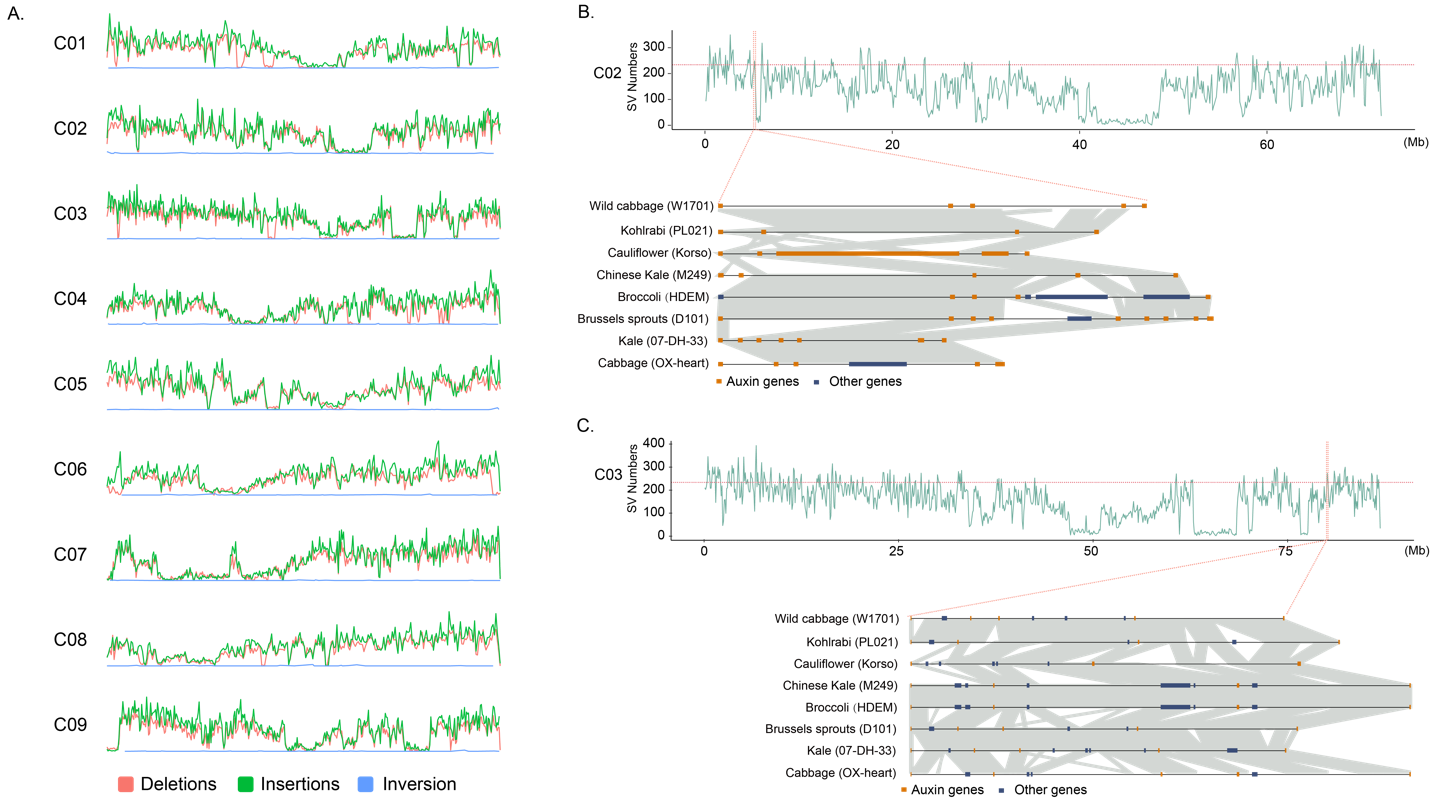

**Supplementary Fig. 5** SV hotspots in the *B. oleracea* genome. (A) Densities of SVs (insertions, deletions, and inversions) across different chromosomes of *B. oleracea*. (B) An SV hotspot on chromosome C02 (5.2-5.4 Mb) harboring 240 SVs associated with five SAUR genes. (C) An SV hotspot on chromosome C03 (80.0-80.2 Mb) harboring 256 SVs associated with five SAUR genes.

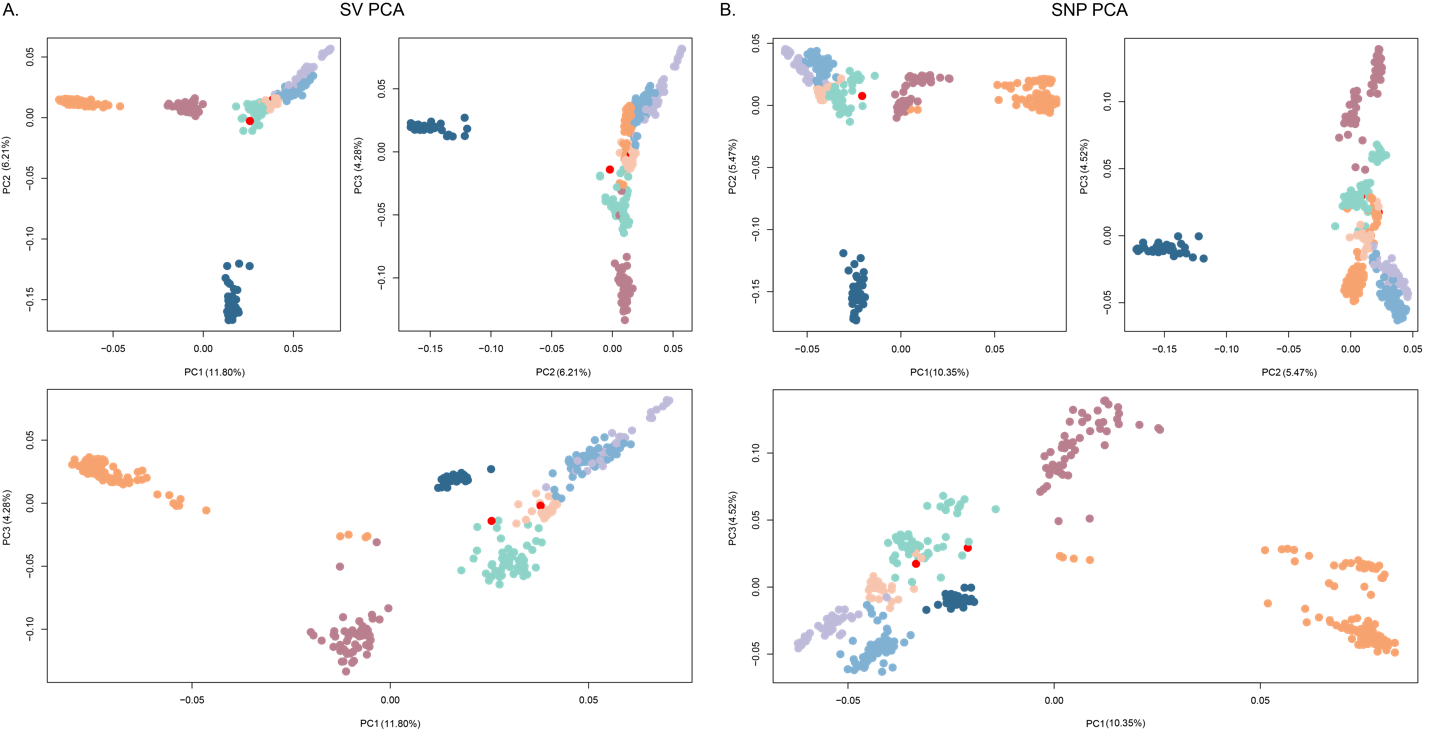

**Supplementary Fig. 6** PCA analyses of 392 *B. oleracea* accessions using SVs (A) and SNPs (B). Colors correspond to different groups: red, wild cabbage; light orange, kale; purple, Brussels sprouts; light blue, cabbage; cyan, kohlrabi; dark blue, Chinese kale; orange, cauliflower; claret, broccoli.

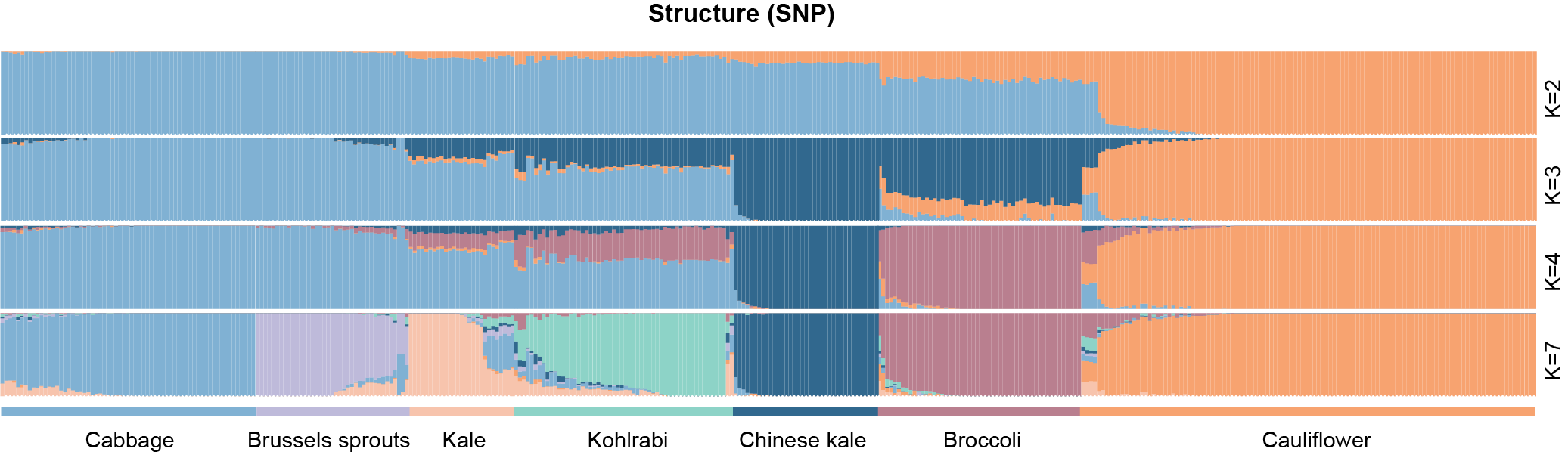

**Supplementary Fig. 7** Model-based clustering of the 392 *B. oleracea* accessions using SNPs with K = 2, 3, 4, and 7.

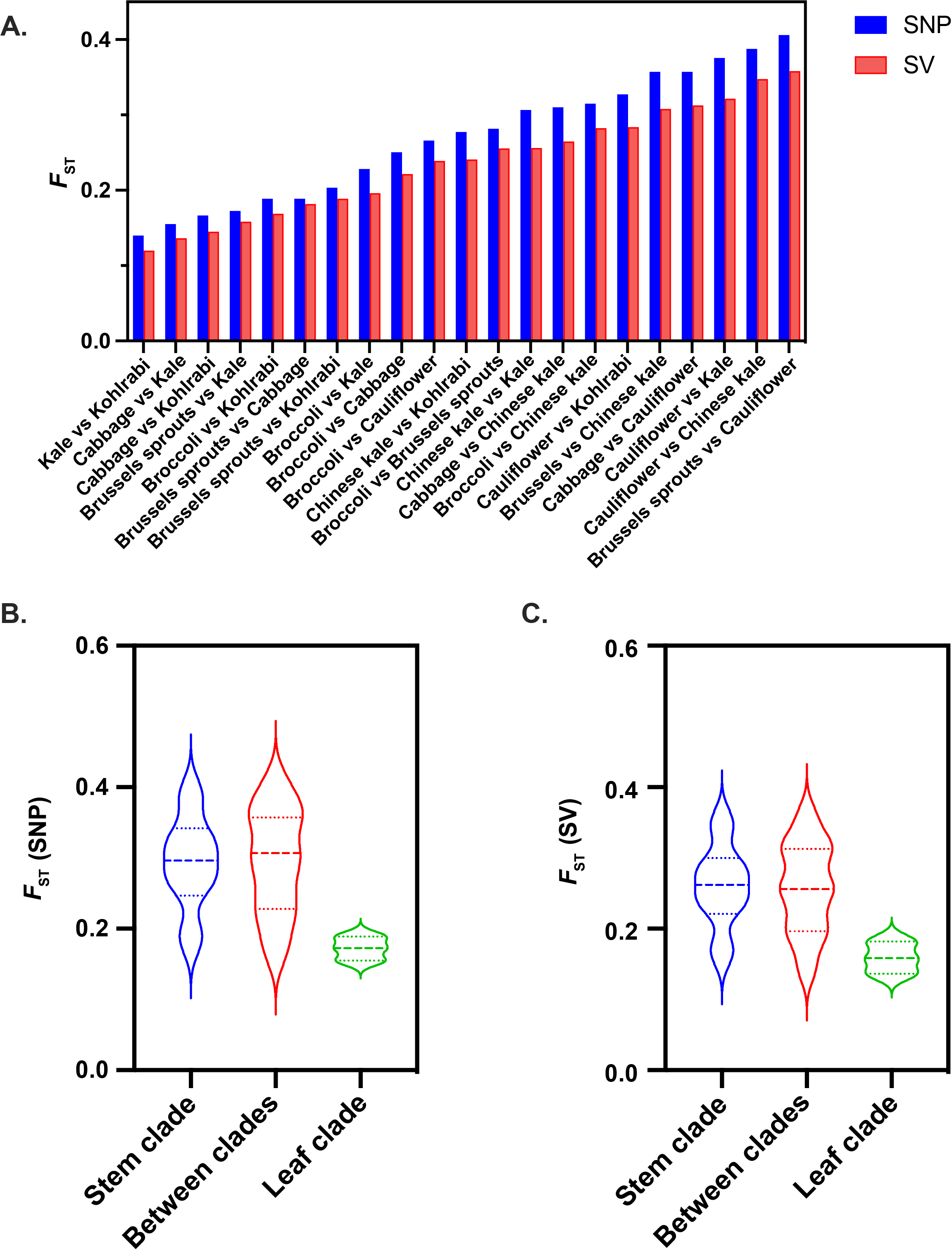

**Supplementary Fig. 8** *F*_ST_ values calculated using SNPs and SVs. (A) *F*_ST_ values of pairwise comparisons between the seven *B. oleracea* morphotypes. (B and C) *F*_ST_ values within the leaf-selected clade, within the stem-selected clade and between the two clades calculated using SNPs (B) and SVs (C).

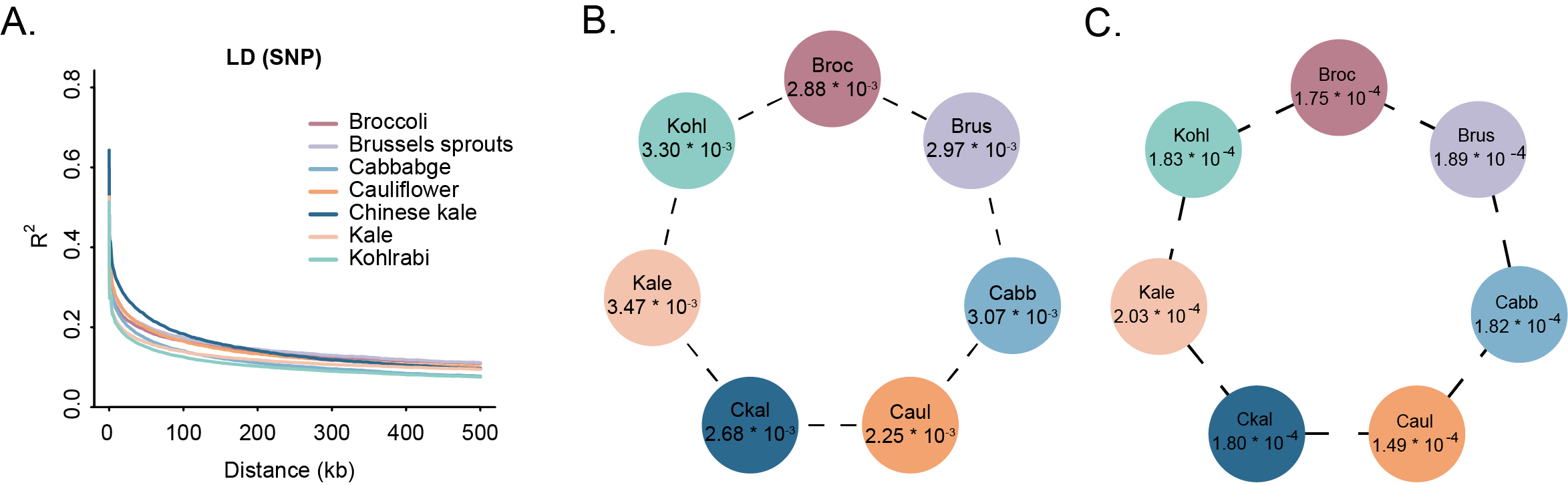

**Supplementary Fig. 9** LD decay and nucleotide diversity (π) of different *B. oleracea* morphotypes. (A) LD decay patterns inferred using SNPs. (B, C) π values of the seven *B. oleracea* morphotypes calculated using SNPs (B) and SVs (C). Broc, broccoli; Brus, Brussels sprouts; Cabb, cabbage; Caul, cauliflower; Ckal, Chinese kale; Kohl, kohlrabi.

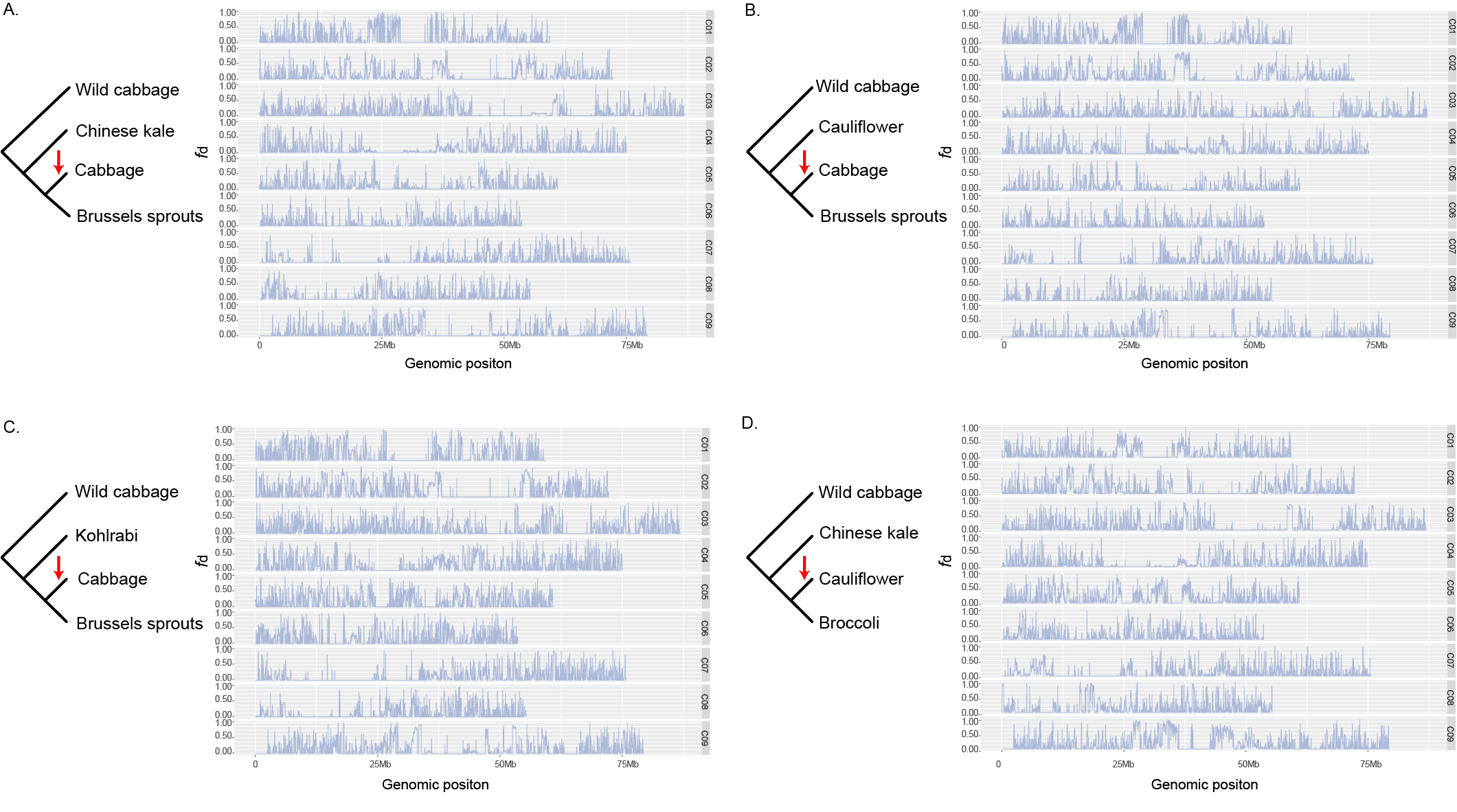

**Supplementary Fig. 10** Gene flows among different *B. oleracea* morphotypes. The four-taxon topology used for modeling introgression in the ABBA–BABA tests identified introgressions from Chinese kale to cabbage (A), from cauliflower to cabbage (B), from kohlrabi to cabbage (C), and from Chinese kale to cauliflower (D) across different chromosomes.

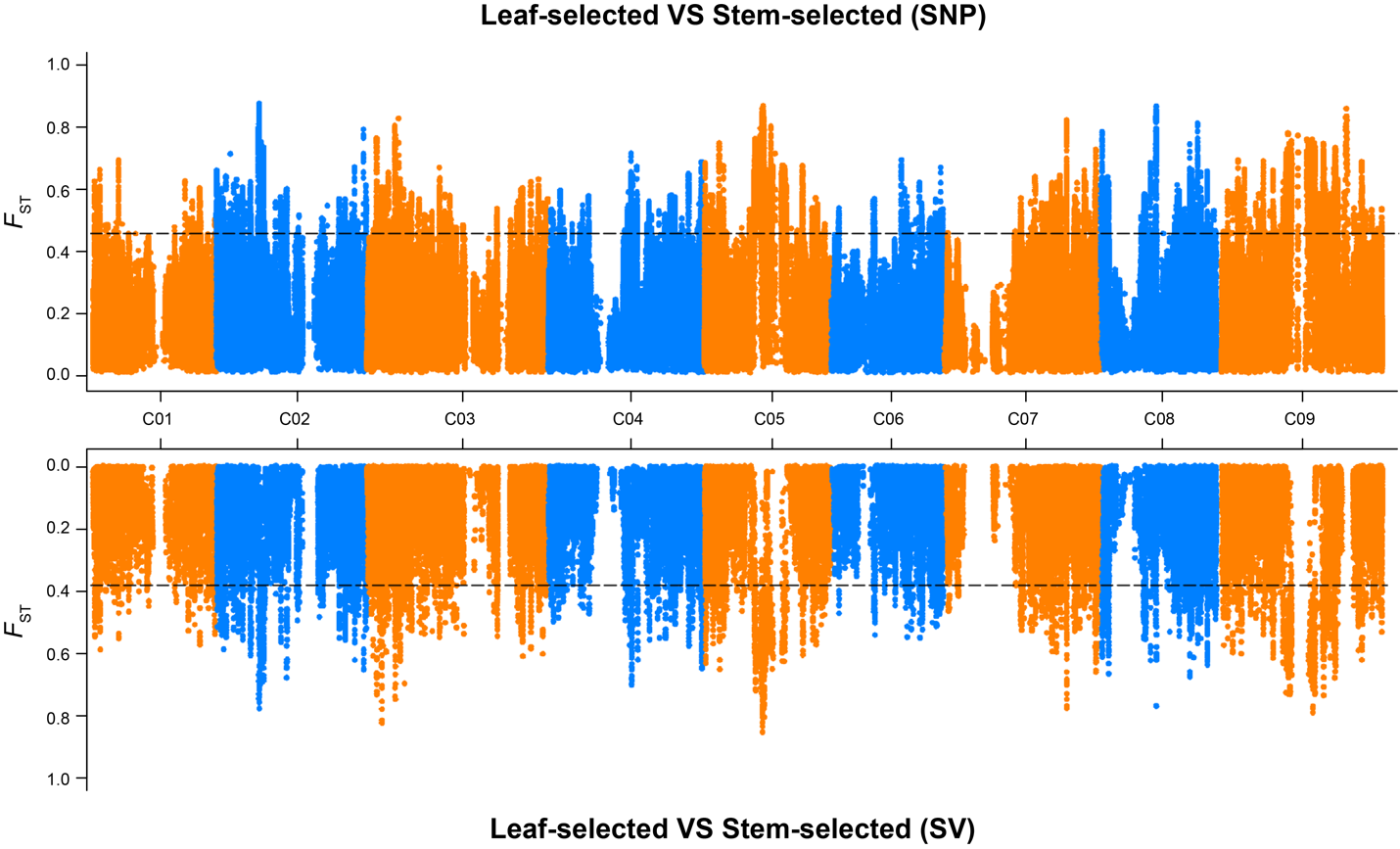

**Supplementary Fig. 11** *F*_ST_ analyses between the leaf- and the stem-selected groups using SVs and SNPs. Dashed horizontal lines indicate the top 5% *F*_ST_.

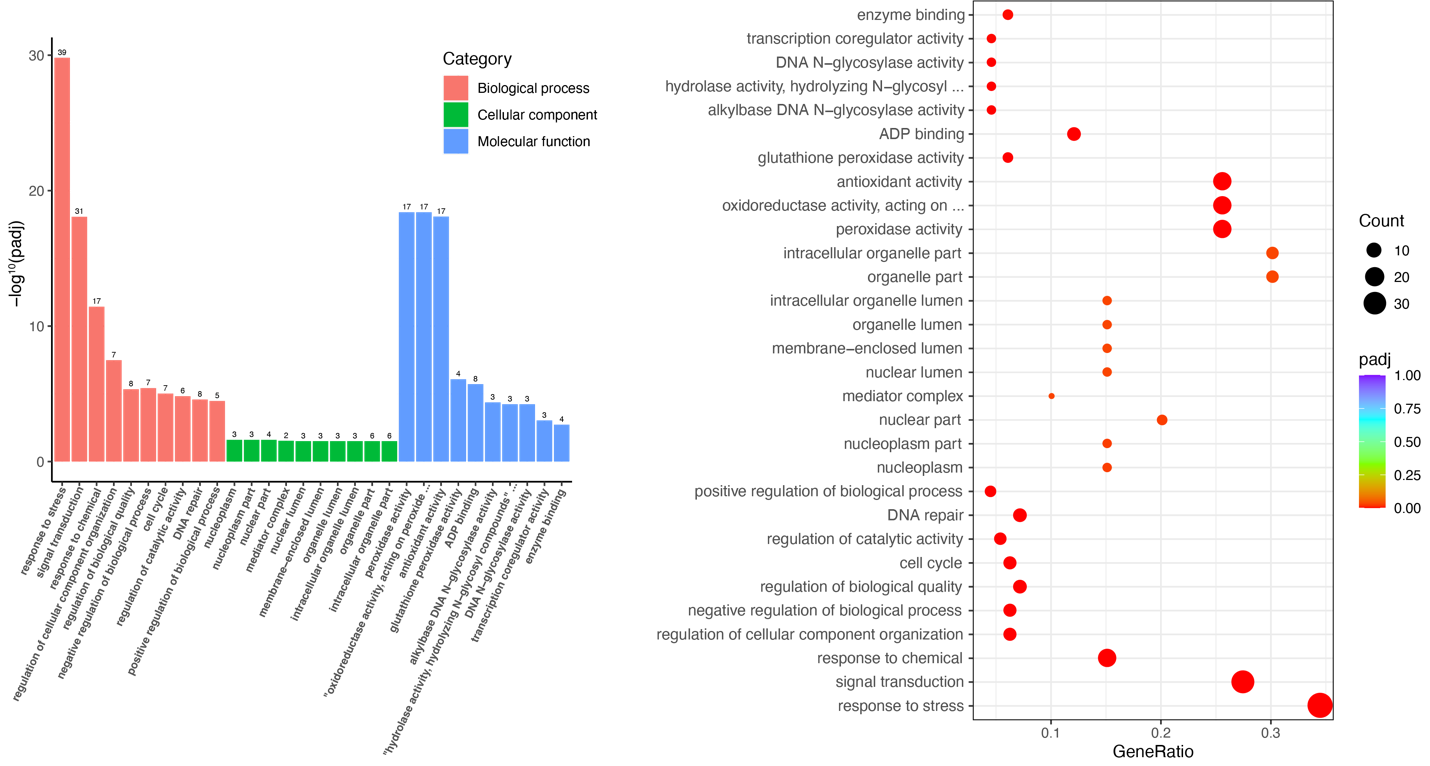

**Supplementary Fig. 12** GO enrichment analysis of selected genes between the leaf- and the stem-selected groups.

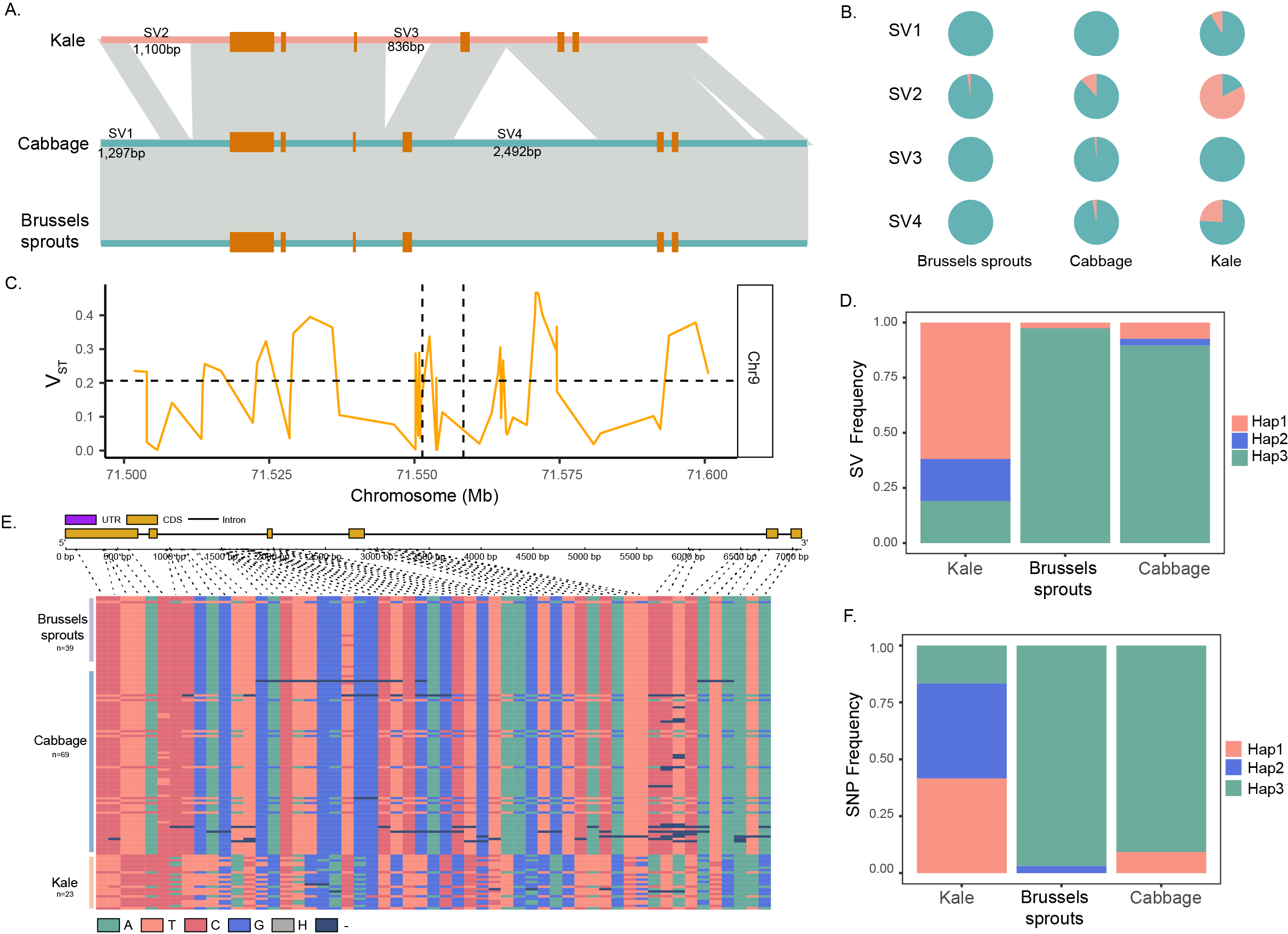

**Supplementary Fig. 13** Structural variations of *BoKAN1.* (A) Four SVs among kale, cabbage, and Brussels sprouts associated with the *BoKAN1* gene. (B) Allele frequencies of the four SVs in kale, cabbage, and Brussels sprouts. (C) Selection of *BoKAN1* inferred by the V_ST_ analysis using SVs. (D) Haplotypes of the four SVs in kale, cabbage, and Brussels sprouts. (E) SNP genotype profiles in the BoKAN1 gene region in kale, cabbage, and Brussels sprouts. (F) SNP haplotypes of *BoKAN1* in kale, cabbage, and Brussels sprouts.

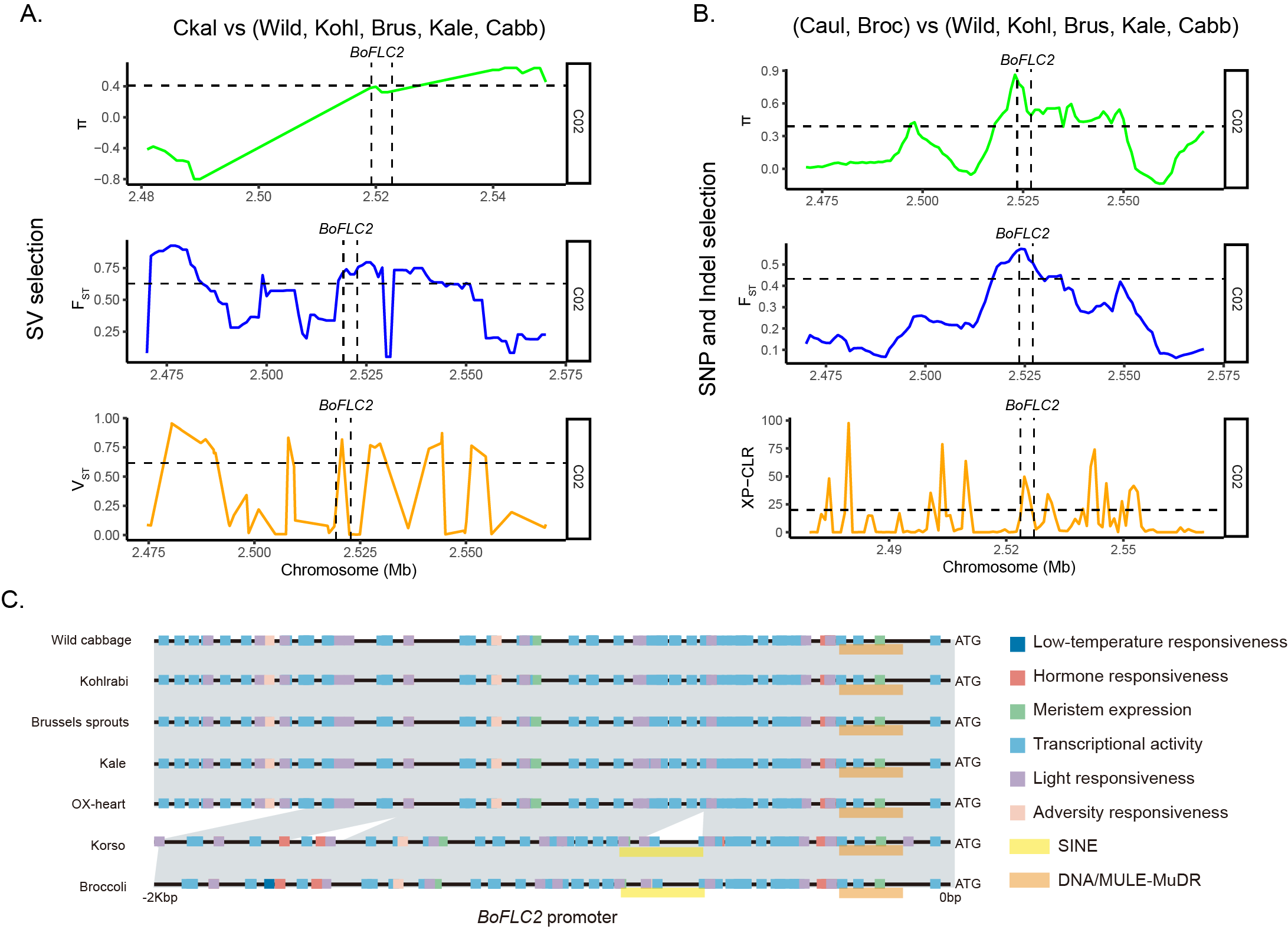

**Supplementary Fig. 14** Selection of the *BoFLC2* gene. (A) Selection signals of *BoFLC2* between Chinese kale and other morphotypes detected by three different methods (π, F_ST_, V_ST_) using SVs. (B) Selection signal of *BoFLC2* between cauliflower plus broccoli and other morphotypes detected using SNPs. (C) Motifs predicted in the promoter region of *BoFLC2* in different morphotypes.

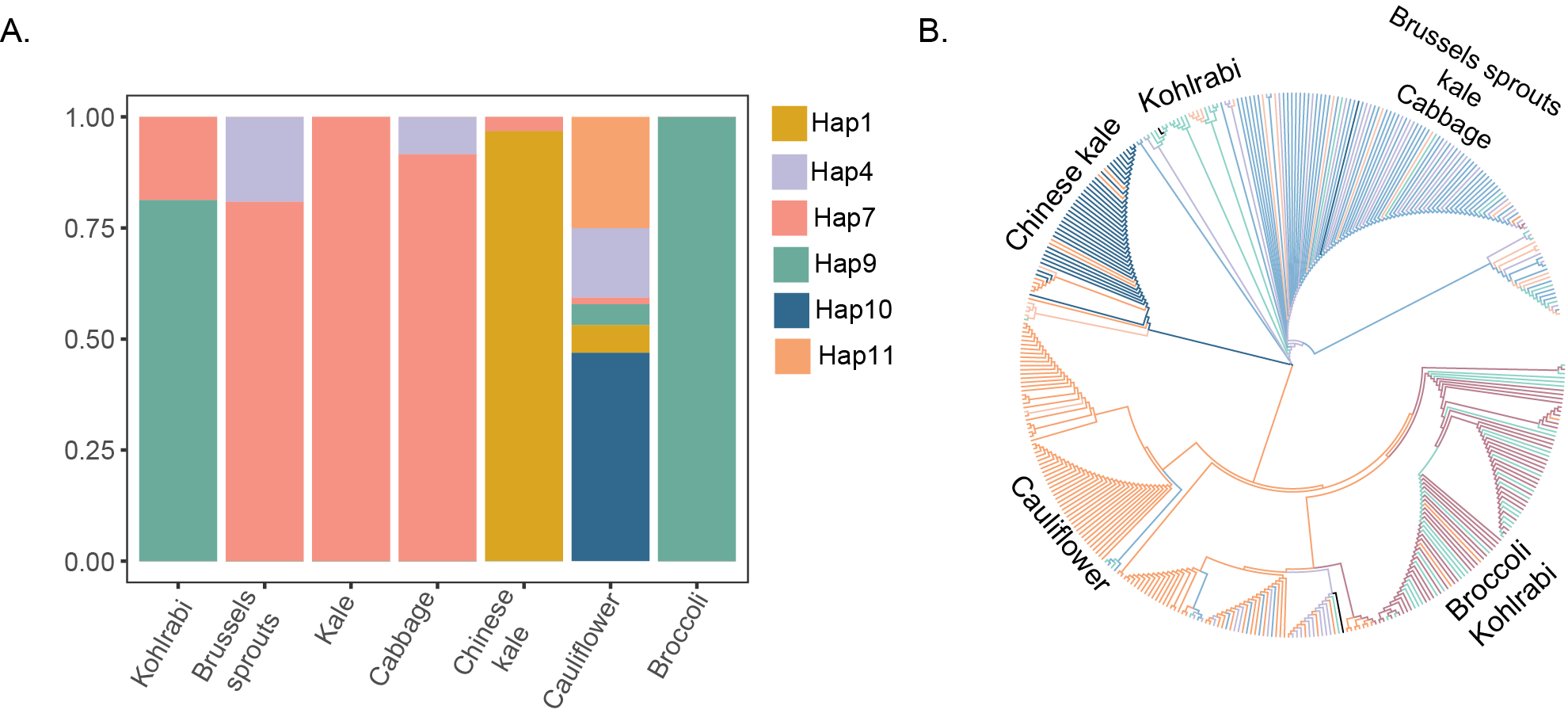

**Supplementary Fig. 15** SNP haplotypes of *BoFLC3*. (A) SNP haplotypes of *BoFLC3* in different morphotypes. (B) Phylogenetic tree of the 392 *B. oleracea* accessions constructed using SNPs in the genomic region of *BoFLC3*.

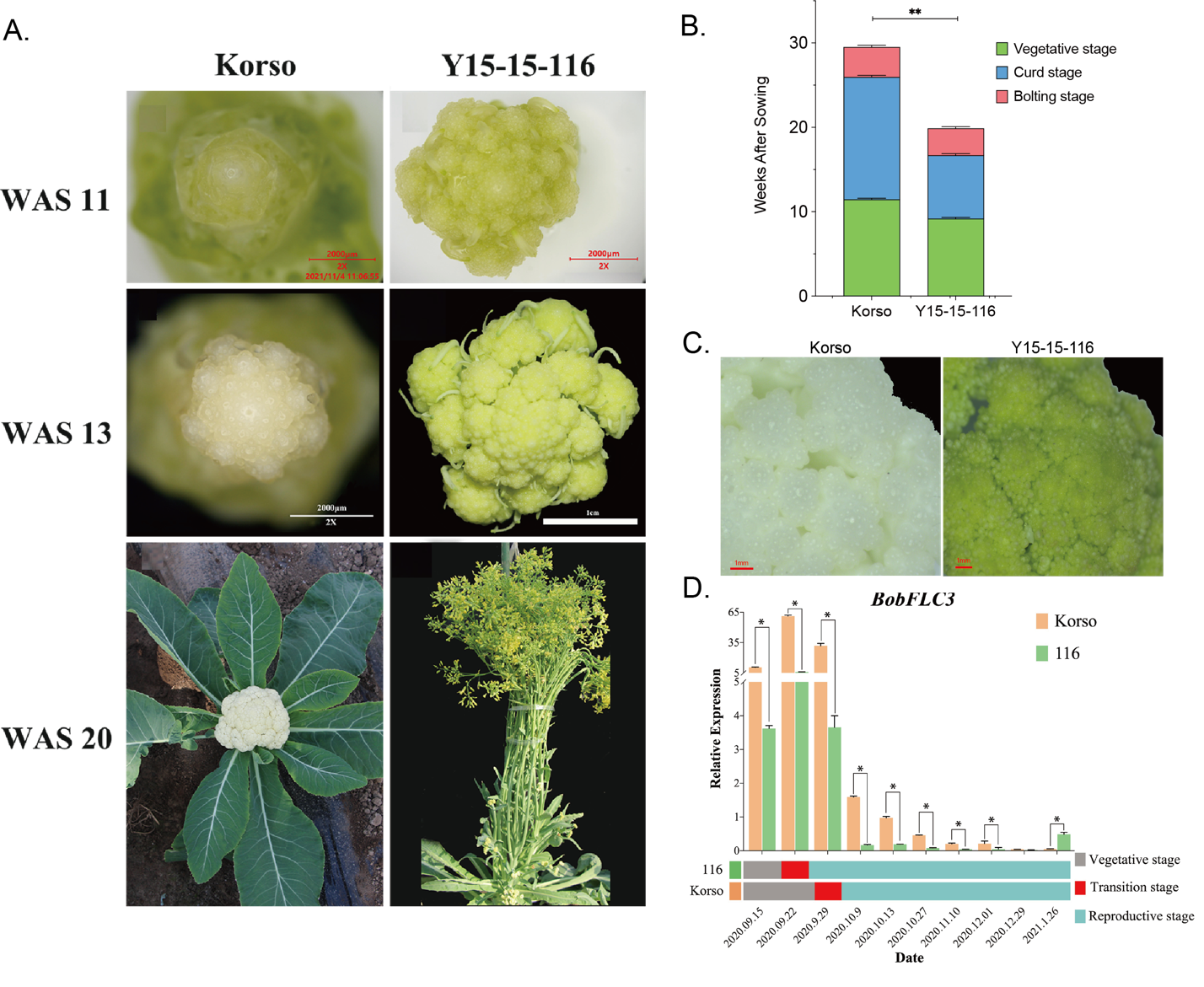

**Supplementary Fig. 16** Phenotype of ‘Korso’ and ‘Y15-15-116’. (A) Curds of ‘Korso’ and ‘Y15-15-116’ at three different developmental stages, 11, 13 and 20 weeks after sowing (WAS). (B) Difference of duration of vegetative stage, curd stage and bolting stage between ‘Korso’ and ‘Y15-15-116’. (C) Curds of ‘Korso’ and ‘Y15-15-116’ at 16 weeks after sowing. (D) Expression of *BoFLC3* in ‘Korso’ and ‘Y15-15-116’ at different developmental stages.
